# A splicing isoform of GPR56 mediates microglial synaptic refinement via phosphatidylserine binding

**DOI:** 10.1101/2020.04.24.059840

**Authors:** Tao Li, Brian Chiou, Casey K Gilman, Rong Luo, Tatsuhiro Koshi, Diankun Yu, Hayeon C Oak, Stefanie Giera, Erin Johnson-Venkatesh, Allie K. Muthukumar, Beth Stevens, Hisashi Umemori, Xianhua Piao

## Abstract

Developmental synaptic remodeling is important for the formation of precise neural circuitry and its disruption has been linked to neurodevelopmental disorders such as autism and schizophrenia. Microglia prune synapses, but integration of this synapse pruning with overlapping and concurrent neurodevelopmental processes remains elusive. Adhesion G protein-coupled receptor ADGRG1/GPR56 controls multiple aspects of brain development in a cell type-specific manner: in neural progenitor cells, GPR56 regulates cortical lamination, whereas in oligodendrocyte progenitor cells, GPR56 controls developmental myelination and myelin repair. Here, we show that microglial GPR56 maintains appropriate synaptic numbers in several brain regions in a time- and circuit-dependent fashion. Phosphatidylserine (PS) on pre-synaptic elements binds GPR56 in a domain-specific manner, and microglia-specific deletion of *Gpr56* leads to increased synapses as a result of reduced microglial engulfment of PS^+^ pre-synaptic inputs. Remarkably, a particular alternatively spliced isoform of GPR56 is selectively required for microglia-mediated synaptic pruning. Our present data provide a ligand- and isoform-specific mechanism underlying microglial GPR56-mediated synapse pruning in the context of complex neurodevelopmental processes.

## Introduction

Microglia, tissue resident macrophages of the central nervous system (CNS), are important for synaptic development, both in promoting synapse formation (Miyamoto et al., 2016; Parkhurst et al., 2013) and in engulfing redundant synapses (Paolicelli et al., 2011; Schafer et al., 2012). Immune molecules such as classical complement components and receptors, CX3CL1/CX3CR1, MHC class I and PirB have been implicated in developmental synaptic refinement (Djurisic et al., 2019; Lee et al., 2014; Paolicelli et al., 2011; Schafer et al., 2012; Stevens et al., 2007; Vainchtein et al., 2018) and in synapse loss in disease models (Bialas et al., 2017; Hong et al., 2016; Sekar et al., 2016; Sellgren et al., 2019; Vasek et al., 2016). Mammalian neurodevelopment involves a succession of overlapping processes beginning with neurogenesis and neuronal migration, which are concurrent with microglial infiltration and morphogenesis. Subsequently, neurite arborization sets the stage for synaptogenesis, circuit establishment, and refinement, as well as myelination. All of these processes entail cell-cell interactions; discovery of molecules involved in multiple processes in a cell-type specific fashion can inform our understanding about how overlapping and sequential programs of intercellular signaling events are coordinately regulated.

The adhesion G protein-coupled receptor (aGPCR) ADGRG1/GPR56 controls several aspects of brain development in a cell type-specific manner by mediating cell-cell and cell-matrix interactions (Langenhan et al., 2016; Singer et al., 2013). During embryonic brain development, GPR56 is expressed in neural progenitor cells and migrating neurons, and interacts with its extracellular matrix (ECM) ligand collagen III to regulate cortical lamination (Jeong et al., 2012a; Jeong et al., 2012b; Luo et al., 2011). In later stages of brain development and throughout postnatal life, GPR56 is highly expressed in glial cells, including astrocytes, oligodendrocyte lineage cells, and microglia (Bennett et al., 2016; Zhang et al., 2014). We recently showed that oligodendrocyte precursor cell (OPC) GPR56 functions together with its microglia-produced ligand, tissue transglutaminase (TG2, gene symbol *Tgm2*), and an ECM component laminin to control developmental myelination and myelin repair (Ackerman et al., 2015; Giera et al., 2015; Giera et al., 2018). Consistent with these findings, germline homozygous loss of function mutations in *GPR56* cause a compound human brain malformation whose phenotype includes aberrant cortical architecture and dysmyelination (Piao et al., 2005; Piao et al., 2004). This phenotype is recapitulated in genetic mouse models indicating conserved GPR56 function (Giera et al., 2015; Li et al., 2008).

Recent studies showed that *Gpr56* is only expressed in yolk sac-derived microglia but not in microglia-like cells engrafted from fetal liver- and bone marrow-derived hematopoietic stem cells, even after long-term adaptation in the CNS *in vivo* (Bennett et al., 2018; Cronk et al., 2018). Furthermore, *Gpr56* expression is promptly lost in primary cultures of microglia (Bohlen et al., 2017; Gosselin et al., 2017). Thus, *Gpr56* is one of few genes that defines the microglial lineage and requires both the appropriate ontogeny and environmental cues for its expression. Motivated by the concept that cell-type-specific functions of GPR56 might coordinate multiple sequential and overlapping neurodevelopmental processes, we tested the hypothesis that microglial GPR56 mediates synapse refinement during postnatal life. Our study results uncover that an alternatively spliced isoform of GPR56 is required for microglia-mediated synapse refinement via binding to phosphatidylserine (PS).

## Results

### PS flags retinal ganglion cell (RGC) synaptic inputs for removal by microglia

How do microglia discriminate which synapse to eliminate? PS is a phospholipid that largely resides on the inner leaflet of the plasma membrane under normal conditions (McLaughlin and Murray, 2005). PS externalization serves as an “eat me” signal for clearance of apoptotic and stressed cells as well as outer segment membranes of retinal photoreceptors (Feng et al., 2002; Neher et al., 2013; Segawa et al., 2011; Tufail et al., 2017). Here, we hypothesize that PS flags synapses for removal. To test this hypothesis, we employed the mouse retinogeniculate system, a classic model to study developmental synaptic pruning (Schafer et al., 2012; Shatz and Kirkwood, 1984). During early postnatal stage, the axons of RGCs extend into the dorsal lateral geniculate nucleus (dLGN) and form excessive synaptic connections with relay neurons. These RGC synaptic inputs undergo pruning by microglia (Schafer et al., 2012). We performed sequential dual labeling with an RGC anterograde tracer - fluorescent cholera toxin B (CTB) and a PS marker - PSVue (Koulov et al., 2003; Smith et al., 2011), a small PS-binding molecule with superior tissue diffusion than pSIVA (an Annexin B12 derivative that binds PS) (Fig. EV1). CTB was intravitreally injected into eyes to trace RGC inputs at P5, and PSVue was injected into the board area between the dLGN and hippocampus at P6 (Fig. 1A). Six hours after PSVue injection, fresh mouse brains were sectioned and imaged under a confocal microscope (Fig. 1B). We observed PSVue colocalizing with some RGC inputs in the dLGN of WT mice (Fig. EV1B). In the developing dLGN, microglial engulfment of synaptic elements peaks at P5 and is dramatically decreased at P10 (Schafer et al., 2012). To investigate whether there are more PS^+^ synapses during the time point of peak synaptic pruning, we performed dual labeling of PSVue and CTB at P6 and P13. Indeed, we observed that nearly 10% of RGC inputs were PSVue-positive at P6, but only 3% were PSVue-positive at P13 (Fig. 1C and D), coinciding with the rise and fall of microglia-mediated synaptic pruning from P5 to P10. To further confirm PS reside with synaptic structures, we performed immunohistochemistry (IHC) on P6 brain for vesicular glutamate transporter 2 (vGlut2), a presynaptic marker specific to RGC inputs in the dLGN (Land et al., 2004), and Homer1, a postsynaptic marker. Indeed, we observed co-localization of PSVue with vGlut2^+^ presynapses and Homer1^+^ postsynapses (Fig. 1E and F).

**Fig. 1.**
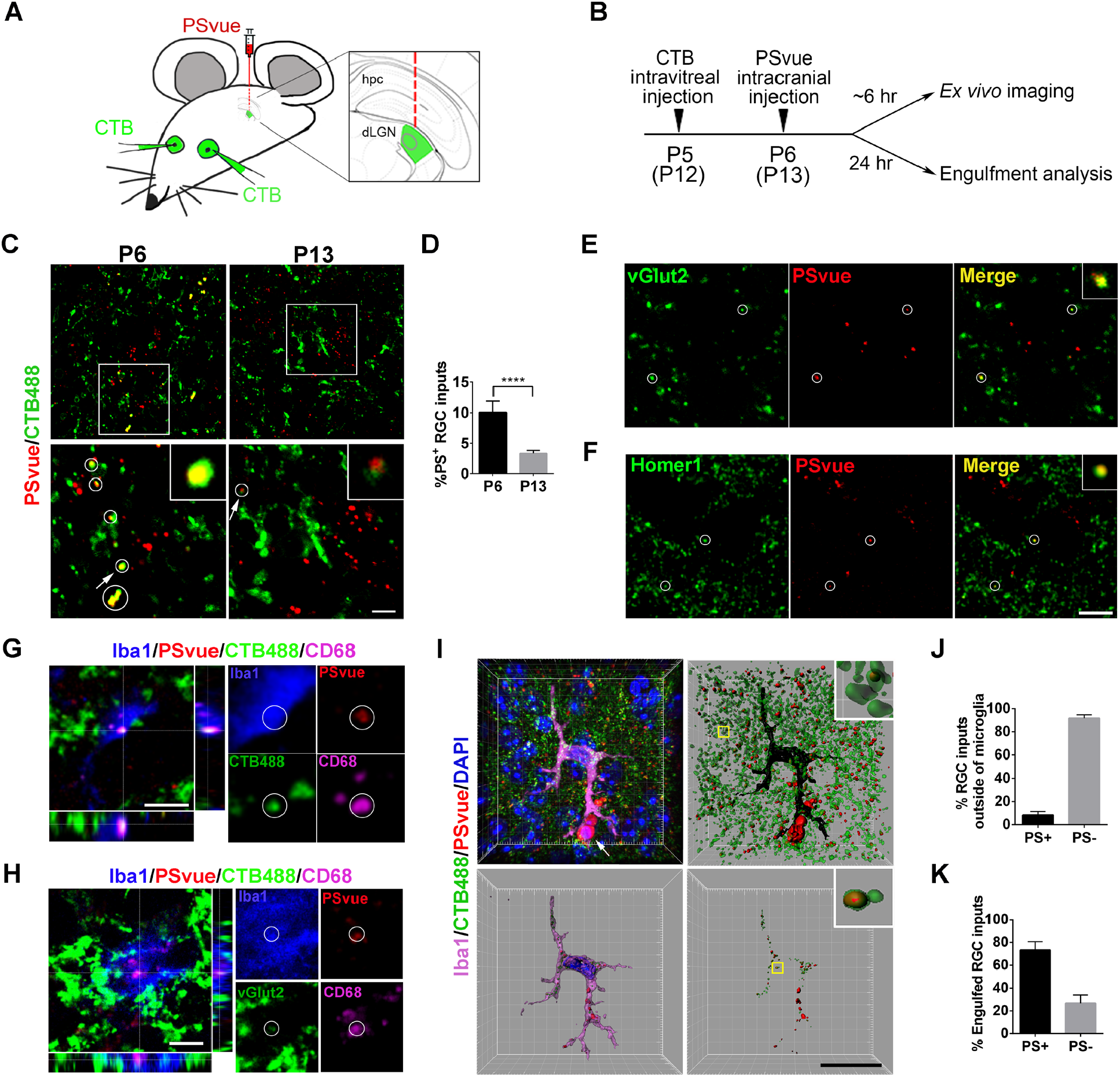
PS flags RGC inputs for removal by microglia during early dLGN development. (A) A schematic drawing of experimental procedure where CTB488 is intraocularly injected, followed by intracranial injection of PSvue550 to dLGN border. (B) A schematic diagram shows the timeline of procedures for *ex vivo* imaging and engulfment analysis. (C) Top panel: Representative images show PS labeling in the WT dLGN at P6 and P13. RGC inputs were labeled with CTB488. Bottom panel: Enlarged images of the boxed region in top panels. Circles indicate PS^+^ RGC inputs. Arrows point to the enlarged PS^+^ RGC inputs in the insets. Scale bar, 5 µm. (D) Quantification of the percentage of PS^+^ RGC inputs in total RGC inputs at P6 and P13. N = 4, ****p < 0.0001. (E) Co-labeling of vGlut2 and PSVue in the dLGN at P6. Circles indicate colocalized vGlut2 and PSVue. (F) Co-labeling of Homer1 and PSVue in the dLGN at P6. Circles indicate colocalized Homer1 and PSVue. Scare bar, 5 µm. (G) Orthogonal sections showed a triple positive PS^+^/CTB488^+^/CD68^+^ RGC inputs inside of microglia. Scare bar, 5 µm. (H) Orthogonal sections showed a triple positive PS^+^/vGlut2^+^/CD68^+^ synapse inside of microglia. Scare bar, 5 µm. (I) A representative image of microglia (upper-left) is surface rendered (bottom-left). The white arrow points to cells which might be apoptotic cells labeled by PSVue. RGC inputs and PSVue outside or inside of microglia are shown in the two right panels. Scare bar, 20 µm. (J) Quantification of the percentage of PS^+^ and PS^-^ RGC inputs outside of microglia in total inputs. N = 4. (K) Quantification of the percentage of engulfed PS^+^ and PS^-^ RGC inputs in total engulfed inputs. N=4. **p < 0.01, ***p < 0.001, ****p < 0.0001, Student’s t-test. Data are presented as mean ± SD.

We next investigated whether microglia engulf PS^+^ RGC synaptic inputs by combining the RGC anterograde tracing, PS labeling, and microglial engulfment assays. RGC inputs were labeled with Alexa 488 conjugated CTB via intravitreal injection at P5, and PS exposed synapses were labeled by PSVue via intracranial injection at P6. Brains were harvested 24 hours later at P7 for analysis of microglial phagocytosis (Fig. 1B). We first tested the hypothesis that microglia engulf PS^+^ pre-synaptic elements rather than free fluorescent dye by performing a negative control experiment using 5-Carboxytetramethylrhodamine (5-TAMRA), the fluorophore component of PSVue that does not bind PS (Hanshaw and Smith, 2005). As shown in Fig. EV2, we observed very sparse 5-TAMRA signals in microglia compared to PSVue, supporting the validity of our assay. In WT animals, we observed that PSVue colocalized with RGC inputs inside microglia (Fig. EV2B). To examine whether RGC inputs were targeted to lysosome in microglia, we did additional staining against CD68, a lysosomal marker for microglia, and detected internalized PS^+^ RGC synaptic inputs colocalized with CD68 (Fig. 1G). Moreover, vGlut2^+^ presynapses were also colocalized with CD68 and PSVue in microglia (Fig. 1H). Next, we quantified the percentage of PS^+^ RGC inputs. We saw less than 10% RGC inputs outside microglia were PS^+^ (Fig. 1I and J), but ∼73% RGC inputs inside microglia were PS^+^ (Fig. 1K), suggesting microglia favorably engulf PS^+^ synaptic inputs, although they do also engulf PS^-^ inputs to a lesser degree (Fig. 1K). Taken together, these data demonstrated that exposed PS flags RGC presynapses for removal by microglia during the early development of dLGN.

### GPR56 binds to phosphatidylserine

*Gpr56* transcript increases in microglia from embryonic stage, and reaches a high level between P3-P6, a period of active microglia-mediated synaptic pruning (Appendix Fig. S1) (Matcovitch-Natan et al., 2016). Considering the facts that *Gpr56* defines true yolk sac-derived microglia (Bennett et al., 2018; Cronk et al., 2018; Matcovitch-Natan et al., 2016) and that BAI1/ADGRB1, another aGPCR family member, recognizes PS (Park et al., 2007), we hypothesized that microglial GPR56 regulates developmental synaptic pruning by binding to PS. GPR56 contains an extensive N-terminal fragment (NTF) followed by a classical seven-transmembrane region and a cytoplasmic tail (Fig. 2A) (Folts et al., 2019). Within the long NTF, there are two functional domains, termed <u>p</u>entraxin/laminin/neurexin/sex-hormone-binding-globulin-like (PLL) and <u>G</u>PCR <u>a</u>utoproteolysis <u>in</u>ducing (GAIN) domains (Araç et al., 2012; Salzman et al., 2016). We engineered recombinant proteins of human immunoglobulin Fc (hFc)-tagged full-length NTF (NTF-hFc) and GAIN-hFc (Fig. 2B).

**Fig. 2.**
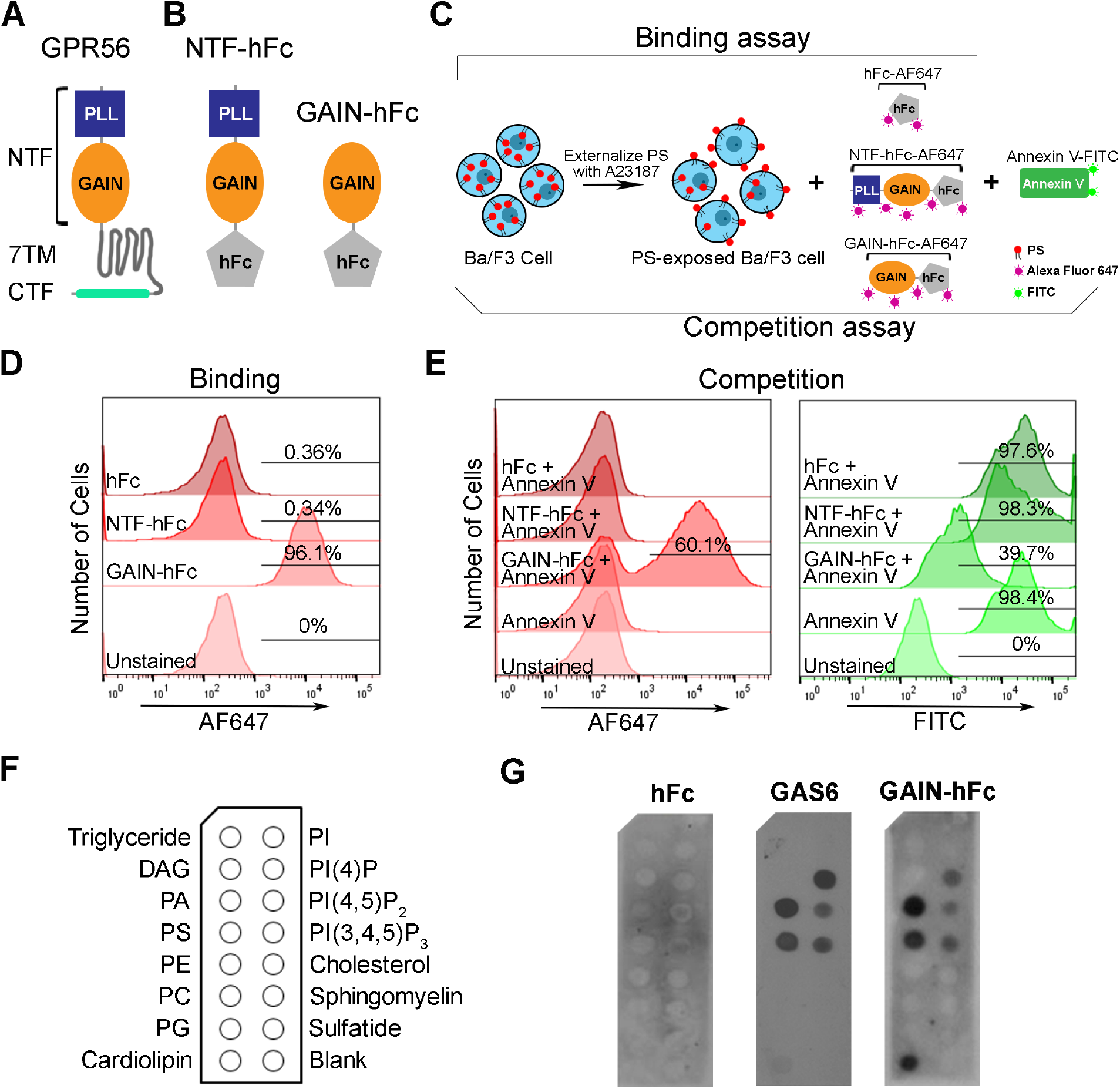
GPR56 binds to PS. (A) A schematic drawing of GPR56 protein structure, with a N-terminal fragment (NTF), a seven transmembrane domain (7-TM) and a C-terminal fragment (CTF). (B) A diagram shows the hFc tag was added to the c-terminal of GPR56-NTF (NTF-hFC) or GAIN domain (GAIN-hFc). (C) A flow chart showing the experimental design of flow cytometry analysis. Briefly, Ba/F3 cells are treated with A23187 to externalize PS. For the binding assay, Alexa Fluor 647-conjugated hFc, GAIN-hFc, or NTF-hFc were incubated with PS-externalized Ba/F3 cells. For the competition assay, Alexa Fluor 647-conjugated hFc, GAIN-hFc, or NTF-hFc were used to compete with FITC-conjugated Annexin V binding. (D) The binding experiment using flow cytometry show that only the GAIN domain binds to PS, similar as Annexin V binding. (E) In the competition experiment, AF647 channel shows that GAIN domain binds 60.1% of the PS^+^ Ba/F3 cells (left). Correspondingly, FITC channel reveals 39.7% Annexin V binding to PS^+^ Ba/F3 cells (right). (F) A diagram of membrane lipid strips spotted with fifteen different lipids. DAG, diacylglycerol; PA, phosphatidic acid; PE, phosphatidylethanolamine; PC, phosphatidylcholine; PG, phosphatidylglycerol; PI, phosphatidylinositol. (G) Direct binding of hFc-tagged GPR56 GAIN proteins to PS and other phospholipids. HFc was used as a negative control, and GAS6 as a positive control.

We investigated whether GPR56 binds PS on cells using flow cytometry analyses of Ba/F3, an IL-3 dependent murine pro-B cell line that externalizes PS upon calcium ionophore A23187 treatment (Appendix Fig. S2) (Suzuki et al., 2010). We first performed a direct binding assay using Alexa Fluor 647 (AF647)-labeled full length NTF, GAIN domain, or hFc (Fig. 2C). FITC-conjugated Annexin V, a known PS-binding protein (Koopman et al., 1994), served as a positive control. Unexpectedly, we found that only the GAIN domain, not the full-length NTF, bound PS (Fig. 2D). To further verify this observation, we performed a competition assay, in which labeled full-length NTF, GAIN domain, or hFc was used to displace Annexin V binding (Fig. 2C). Indeed, we confirmed that the GAIN domain, but not full length NTF, competed with Annexin V for binding to PS (Fig. 2E). The PLL and GAIN domains are constrained by an interdomain disulfide bond at two cysteine residues C121 and C177 (Salzman et al., 2016). It is conceivable that the PLL domain blocks GAIN domain binding to PS in the full-length NTF.

To test whether GPR56 GAIN domain binds to other phospholipids besides PS, we performed a protein-lipid overlay experiment using Membrane Lipid Strips, as previously described (Park et al., 2007). GAS6, a known PS-binding protein, was used as a positive control. Interestingly, we found GAIN domain has a similar lipid-binding profile as GAS6, except cardiolipin (Fig. 2F and G). In this *in vitro* membrane-based assay, GPR56 GAIN domain was able to bind phosphatidic acid (PA), phosphatidylserine (PS), cardiolipin, PI(4)P, PI(4,5)P_2_ and PI(3,4,5)P_3_. Given that PA, PI(4)P, PI(4,5)P_2_ and PI(3,4,5)P_3_ are normally present on the inner leaflet of the cell plasma membrane (Ingolfsson et al., 2014), and cardiolipin is a phospholipid almost exclusively located in the inner mitochondrial membrane (Paradies et al., 2014), GAIN domain binding to those lipids carries unclear biological relevance in the context of our current study.

### GPR56 S4 variant is dispensable for cortical development and CNS myelination but is essential for microglia-mediated synaptic refinement

S4 is an alternatively spliced GPR56 isoform that initiates at an alternative ATG start codon in exon 4, resulting in a GPR56 variant that contains only the GAIN domain in its extracellular region, in both humans and mice (Fig. 3A) (Kim et al., 2010; Salzman et al., 2016). We published the cortical phenotype of the germline *Gpr56* gene-targeted mice (Li et al., 2008), before the discovery of *Gpr56* splicing variants. At that time, our data suggested that this line represented a null allele in regard to cortical phenotype. Subsequently, it was characterized as a hypomorphic allele, represented by selective expression of the *Gpr56 S4* variant (Fig. 3B, Appendix Fig. S3) (Salzman et al., 2016). In the present report, this genetic model is referred to as *Gpr56 S4* (Fig. 3C). To extend our investigation of *Gpr56* biology, we deleted *Gpr56* exons 4-6, yielding a global null mutant termed as *Gpr56 null*, lacking both full length GPR56 and its S4 variant (Fig. 3C). Based on the fact that the PLL domain binds collagen III (Luo et al., 2012), which is the relevant GPR56 ligand in the developing cerebral cortex, we speculated that the S4 isoform would not be required for cerebral cortical lamination. Indeed, we observed a comparable cortical ectopia size and distribution between *Gpr56 S4* and *Gpr56 null* mice (Fig. 3D-F). GPR56 also drives myelination and myelin repair via the PLL domain and TG2 interaction (Giera et al., 2015; Giera et al., 2018; Singer et al., 2013). As we expected, we observed comparable decreased myelination in *Gpr56 null* and *Gpr56 S4* mice (Fig. 3G and H), which suggested that the GPR56 S4 isoform is dispensable in OPC GPR56 regulation of CNS myelination.

**Fig. 3.**
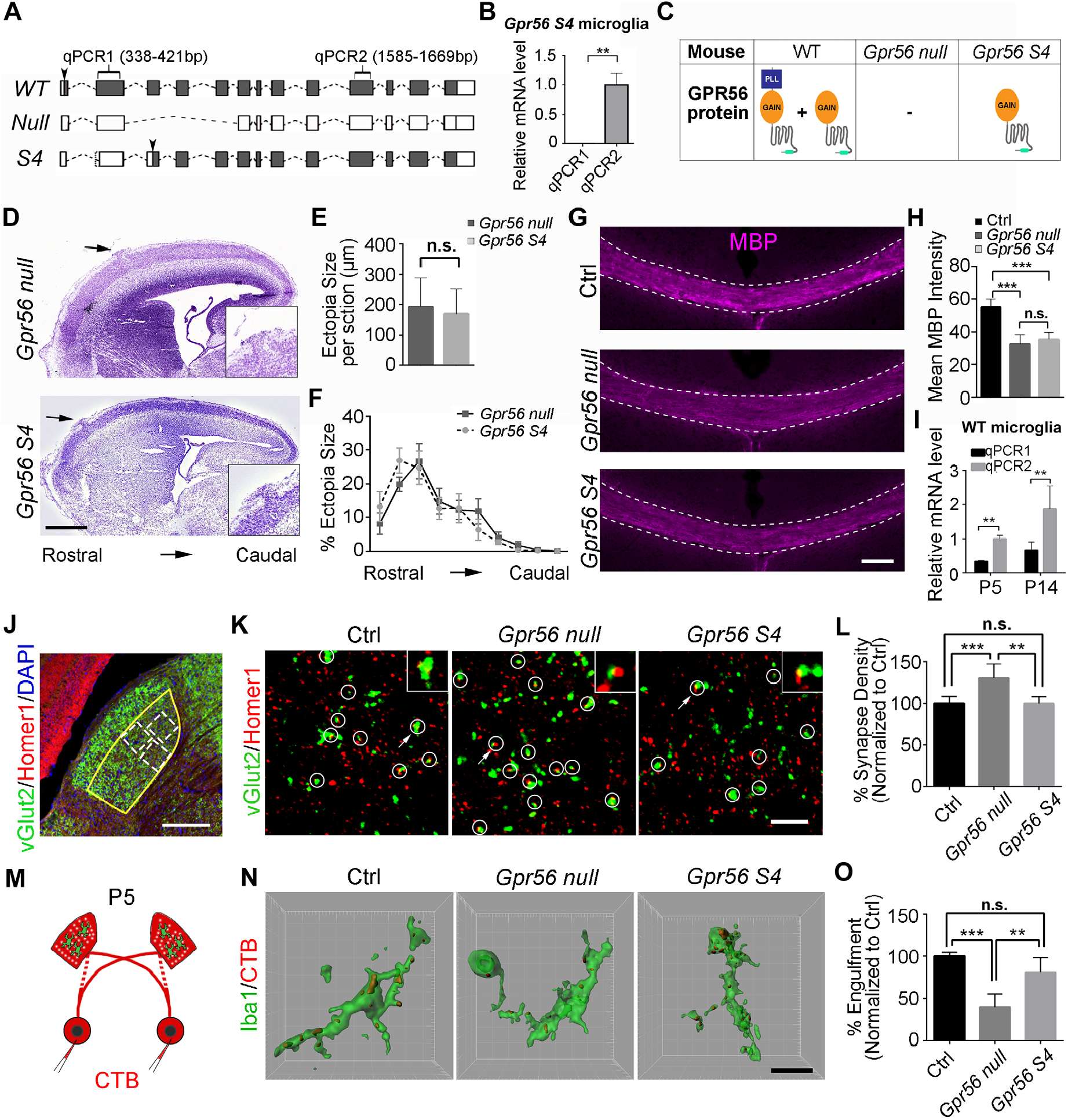
GPR56 S4 variant is required for synaptic refinement in the dLGN. (A) A diagram showing *Gpr56 WT, null* and *S4* transcripts. Solid boxes indicating exons that are transcribed. Arrowheads indicate the translation start sites. (B) *Gpr56* transcript levels in isolated microglia from *Gpr56 S4* mice. N = 3, **p < 0.01 by Student’s t-test. (C) A table showing the expression status of *Gpr56* WT and S4 variants in microglia of different transgenic mice. (D) Representative images of Nissl staining of *Gpr56 null* and *Gpr56 S4* E16.5 neocortex. Arrows indicate cortical ectopias that are shown in insets. (E) Quantification of ectopia size per section. N = 6, p = 0.64 by Student’s t-test. (F) The distribution of ectopia from rostral to caudal cortex. N = 3, F (9, 81) = 0.65, p = 0.75 by two-way ANOVA with Bonferroni’s post hoc test. (G). Myelin basic protein (MBP) staining of corpus callosum in P28 controls, *Gpr56 null* and *Gpr56 S4* mice. Scale bar, 100 µm. (H) Quantification of MBP intensity in corpus callosum. N = 5, ***p < 0.001 by one-way ANOVA with Tukey’s post-hoc test. (I) *Gpr56* transcript levels in WT microglia at P5 and P14. N = 4, **p < 0.01, two-way ANOVA with Sidak’s multiple comparisons test. (J) An overview of vGlut2 and Homer1 staining in dLGN at P10. The yellow outline indicates the dLGN core, and the dotted boxes show where synapses are quantified. Scale bar, 200 µm. (K) Representative images of vGlut2/Homer1 staining in the dLGN of control, *Gpr56 null*, and *Gpr56 S4* brains at P10. Arrows pointing to the enlarged synapse in the insets. Scale bar, 5µm. (L) Relative vGlut2/Homer1 synapse density in dLGN. N=10 (Ctrl), N=6 (*Gpr56 null*), N=3 (*Gpr56 S4*). **p < 0.01, *** p < 0.001, one-way ANOVA with Tukey’s post-hoc test. (M) A schematic representation of the *in vivo* engulfment assay. CTB are injected into both eyes at P4, and anterogradely trace RGC projections to the dLGN. After 24 hours, the brains are collected and analyzed at P5. (N) Representative images and surface rendered microglia (green) in which CTB^+^(red) RGC inputs were engulfed. Scale bar, 10µm. (O) Quantification of the percentage of engulfed RGC inputs in controls, *Gpr56 null* and *Gpr56 S4* microglia. N = 4, **p < 0.01, *** p < 0.001, one-way ANOVA with Tukey’s post-hoc test. Data are presented as mean ± SEM in (F). For other statistics analysis, data are presented as mean ± SD.

Based on PS binding data (Fig. 2), we hypothesized that GPR56 S4 would be required for microglia-mediated synaptic pruning. Supporting this hypothesis, we found that *Gpr56 S4* is the major transcript in microglia as determined by qPCR analysis of microglia isolated from P5 and P14 wild type (WT) mouse brains (Fig. 3I). To test whether GPR56 S4 is implicated in synaptic refinement, we first examined synaptic density in the dLGN (Schafer et al., 2012). To visualize retinogeniculate synaptic connections, we performed double immunohistochemistry (IHC) for vGlut2 and Homer1, followed by confocal imaging in the dLGN core region (Fig. 3J), to resolve juxtaposed pre- and post-synaptic structures. Colocalized vGlut2 and Homer1 puncta (vGlut2^+^/Homer1^+^) were counted as retinogeniculate synapses. Strikingly, germline *Gpr56 null* exhibited significantly increased synapse density at P10, while *Gpr56 S4* and controls showed comparable synapse density (Fig. 3K and L). Interestingly, we observed significantly increased vGlut2^+^ presynaptic signals, but not Homer1^+^ postsynaptic inputs (Appendix Fig. S4). Taken together, our results support that the GPR56 S4 isoform is crucial to synapse development.

To delineate whether this change resulted from a reduction in microglial engulfment of synapses, we performed *in vivo* engulfment assays as previously described (Schafer et al., 2012; Vainchtein et al., 2018). Mice received intraocular injection of CTB at P4 and were sacrificed 24 hours later for analysis, as peak pruning occurs around P5 in the murine retinogeniculate system (Fig. 3M). Indeed, *Gpr56 null* microglia exhibited significantly reduced engulfment of RGC inputs, while *Gpr56 S4* microglia showed a comparable engulfment ability as controls (Fig. 3N and O). Taken together, these results support the concept that the GPR56 S4 isoform is essential to microglia-mediated synaptic pruning.

### Deleting microglial *Gpr56* results in excess synapses in the dLGN during postnatal development

To specifically investigate the function of microglial GPR56 in synaptic refinement, we generated microglia-specific *Gpr56* conditional knockout mice by crossing mice harboring a conditional *Gpr56*^*fl/fl*^ allele (Giera et al., 2015) with *Cx3cr1-Cre* transgenic mice (Yona et al., 2013). Since *Cx3cr1-Cre* is a direct knock-in transgenic mouse line, we used *Gpr56*^*fl/fl*^; *Cx3cr1-Cre*^*+/-*^ mice as conditional knockouts (CKO) and *Gpr56*^*+/+*^; *Cx3cr1-Cre*^*+/-*^ as controls. Western blot and qPCR experiments did not detect GPR56 protein or *Gpr56* transcript in microglia isolated from CKOs and their controls (Fig. EV3A and B). We further performed RNAscope analysis for *Gpr56* in the prefrontal cortex of P30 mice (Fig. 4A). We detected minimal, if any, *Gpr56* transcripts in *Gpr56 null* and CKO microglia, comparing to controls (Fig. 4B). In contrast, *Gpr56* transcripts were abundant in non-microglial cells in both CKO and controls (Fig. 4B), confirming *Gpr56* was cell-specifically deleted in microglia. To further confirm the specificity of Cre-Lox recombination in microglia under the *Cx3cr1-Cre* driver, we crossed *Cx3cr1-Cre* with a reporter line *Rosa-GFP*^*fl*^ and demonstrated restricted GFP expression in microglia (Appendix Fig. S5). To investigate whether deleting microglial *Gpr56* affected microglial cellular properties, we asked whether there were differences between control and CKO in microglial cell density, coverage, morphology, and amounts of phagocytic machinery (defined as %CD68^+^ cells) in the dLGN of P5 mice. This comprehensive analysis showed no significant differences (Fig. EV3C-J), enabling a functional evaluation of microglial GPR56 in synapse development.

**Fig. 4.**
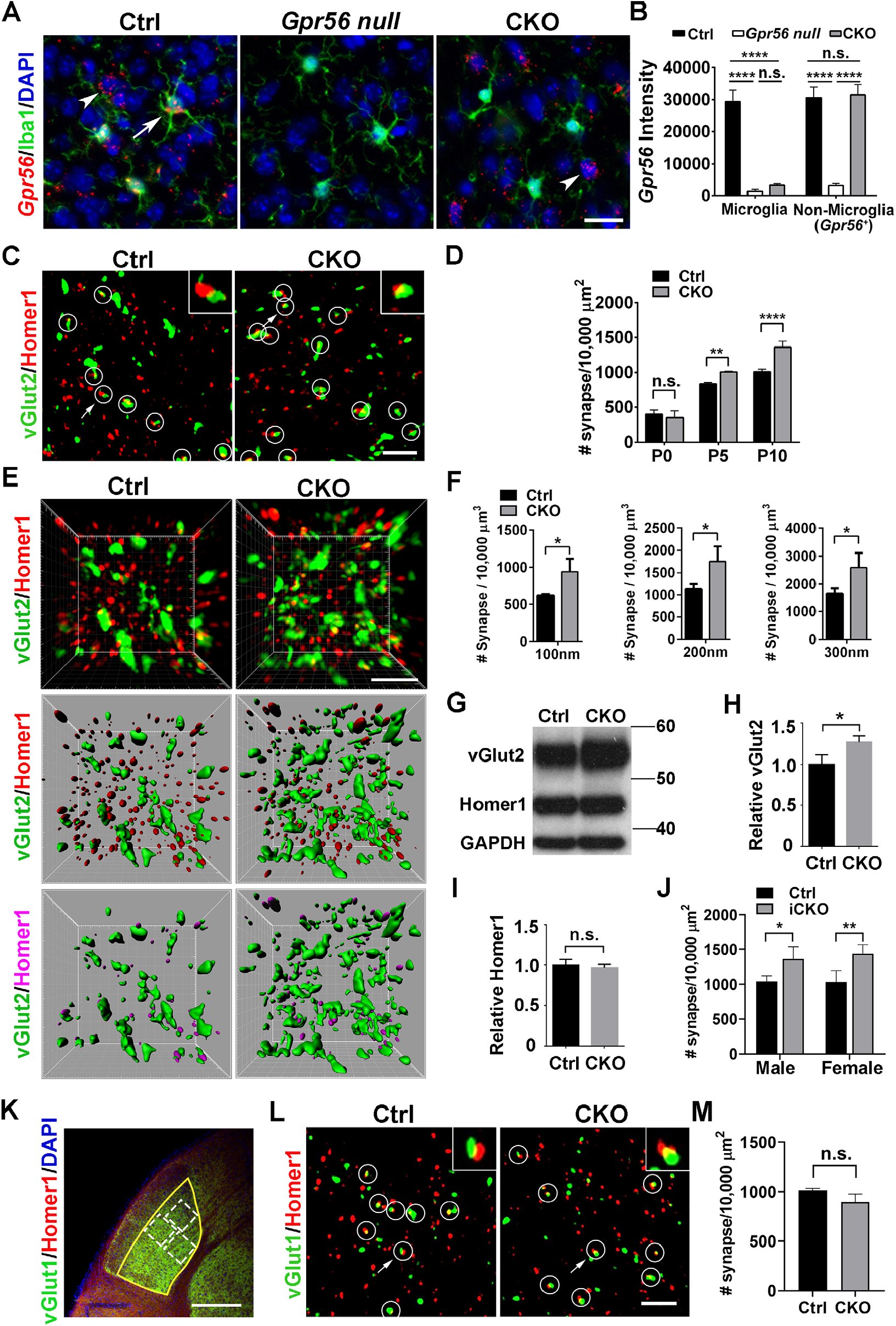
Microglial GPR56 is required for synaptic development in the dLGN. (A) *Gpr56* is deleted in CKO microglia. RNAscope shows *Gpr56* transcripts co-localize with microglia (arrow) in controls, but not in CKO, and is absent in all cell types in the global *Gpr56* KO (*Gpr56 null*). Arrowhead indicates non-microglia cells expressing *Gpr56*. RNAscope was performed in the prefrontal cortex of P30 mice. Scale bar, 20 µm. (B) Quantification of *Gpr56* fluorescence signals in (A). N = 20-50 cells for each. (C) Confocal images of vGlut2 (green) labeling retinal ganglion cell (RGC) presynaptic terminals and Homer1 (red) for post-synapses in the dLGN of CKO and controls at P10. Overlapped vGlut2 and Homer1 are quantified as synapses, indicated by white circles. Arrows pointing to the synapse enlarged in the upper right-hand inset. Scale bar, 5 µm. (D) A time course of dLGN synapse density (vGlut2^+^/Homer1^+^) between WT and controls. N = 3 for P0, N = 4 for P5, N=3 for P10. (E) Representative super-resolution images and 3D-reconstructed images of vGlut2 and Homer1 staining of P8 dLGN. Green objects represent surface rendering of vGlut2^+^ presynaptic terminals. Red spots are rendered Homer1^+^ post-synapses. Magenta spots represent Homer1^+^ post-synapses within a distance of 300nm from the nearest vGlut2^+^ surface. Scale bar, 2 µm. (F) Quantification of Homer1^+^ spots adjacent to vGlut2 surface within 100nm, 200nm, and 300nm, respectively. N = 3. (G) Western blot of vGlut2 and Homer1 using microdissected dLGN tissue from P8 mice. (H, I) Quantification of vGlut2 and Homer1 expression in CKOs and controls. N = 3, *p = 0.027. (J) Quantification of synapse density (vGlut2^+^/Homer1^+^) in the dLGN of both male and female iCKO mice at P10. N = 3 for male, N = 4 for female. (K) An overview of vGlut1 and Homer1 staining in dLGN at P10. The yellow outline indicates the dLGN core, and the dotted boxes show where synapses are quantified. Scale bar, 200 µm. (L) Representative images of vGlut1^+^ (green) presynaptic terminals and Homer1^+^ (red) postsynapses in the dLGN of CKO and controls at P10. Arrows pointing to the synapse enlarged in inset. Scale bar, 5 µm. (M) Quantification of synapse density (vGlut1/homer1) at P10 in the dLGN of CKO mice and controls. N=3 (WT), N=5 (CKO), p = 0.053. *p < 0.05, **p < 0.01, ****p < 0.0001 using Student’s t-test (F, H, I and M) or two-way ANOVA followed by Bonferroni’s post hoc test (B, D and J). Data are presented as mean ± SD.

We first conducted a time course study of synaptic density from P0 to P10 in the dLGN. At birth (P0), we observed comparable retinogeniculate synapse density (vGlut2^+^/Homer1^+^) between CKO mice and their controls. At P5 and P10, we observed a steady increase in synaptic density by ∼21% and 35%, respectively, in CKO mice compared to their age- and sex-matched controls (Fig. 4C and D, Appendix Fig. S6). To further confirm this phenotype, we applied structured illumination microscopy (SIM) to resolve juxtaposed pre- and post-synaptic structures with a higher resolution (Hong et al., 2016; Hong et al., 2017). Previous transmission electron microscopy (EM) studies estimated a distance of ∼ 80nm between Homer1 and the presynaptic cleft (Dani et al., 2010) and a span from 0 - 200nm between the vGlut2 vesicles and the pre-synaptic cleft (Fujiyama et al., 2004). In our analysis of 3D reconstructions of the SIM images, we defined a synapse when the distance from the center of a Homer1 immunoreactive spot to the nearest vGlut2 surface was between 0 - 300 nm (Movie. EV1). Using this approach, we revealed that CKO mice had significantly increased synapse numbers compared to controls at P8 (Fig. 4E and F, Appendix Fig. S7). Importantly, SIM and confocal analyses yielded consistent synaptic changes (Fig. 4D and F), supporting the usage of confocal imaging and analyses for other brain regions and transgenic mice.

Consistent with increased vGlut2^+^/Homer1^+^ synapse numbers in CKOs, western blot analysis of microdissected dLGN from P8 animals showed increased vGlut2 protein levels in CKOs as compared to control brains (Fig. 4G and H), but no change in Homer1 protein levels (Fig. 4I). It is worth noting that the increased synapse density was not the result of altered RGC number, since CKO had comparable brain-specific homeobox/POU domain protein 3A (Brn3a, a marker of RGC (Nadal-Nicolas et al., 2009)) positive RGC density compared to controls (Fig. EV4). Taken together, our data suggested that microglial GPR56 is important for normal synaptic refinement in dLGN.

To address whether the observed synaptic density increase was a consequence of a postnatal event, we generated animals that enabled inducible deletion of microglial *Gpr56* by crossing *Gpr56*^*fl/fl*^ mice with *Cx3cr1-CreER* mouse line postnatally (Yona et al., 2013). *Gpr56*^*fl/fl*^; *Cx3cr1-CreER*^*+/-*^ mice were used as inducible conditional knockouts (iCKO), and *Gpr56*^*+/+*^; *Cx3cr1-CreER*^*+/-*^ mice were used as controls. Tamoxifen was administered to both iCKO and controls at P1-P3 and brains were analyzed at P10. We observed comparable increases in retinogeniculate synaptic density in both male and female iCKO mice, in comparison to their age-matched controls, indicating that there is no sexual dimorphism in microglial GPR56 function in the context of synaptic development (Fig. 4J). Furthermore, iCKO and CKO mice showed a quantitatively equivalent synapse phenotype at P10 (Fig. 4D and J), indicating the observed synaptic phenotype is a postnatal event. Additionally, this result demonstrated that it is appropriate to use CKO mice for most of the remaining studies.

The dLGN also receives vGlut1^+^ projections from neocortical layer VI (Fitzpatrick et al., 1994; Fujiyama et al., 2003; Thomson, 2010), which act as modulatory inputs that flexibly tune postsynaptic activity in target cells (Sherman and Guillery, 1996, 1998). In contrast to increased vGlut2^+^ retinogeniculate synapses, there were no significant changes in the density of vGlut1^+^ corticogeniculate synapses at P10 (Fig. 4K-M), indicating that microglial GPR56 functions in a synapse-specific manner and does not modulate neocortical inputs in dLGN during the developmental stages investigated. Taken together, these findings demonstrate that microglial GPR56 is necessary for retinogeniculate synaptic development in early dLGN.

### Microglial GPR56 regulates hippocampal synaptic development

To address whether microglial GPR56 affects synaptic development in other brain regions, we examined synaptic density in the hippocampus at P10 and P21. In the hippocampus, Schaffer-collateral axons from CA3 synapse on CA1 pyramidal neurons exclusively on dendritic domains in the stratum (S.) oriens and S. radiatum (Swanson et al., 1978). Perforant-path axons from the entorhinal cortex form synapses on CA1 region pyramidal neurons exclusively on dendritic domains in the S. lacunosum-moleculare, a critical neural circuit for temporal associative memory (Suh et al., 2011). We found increased synapse densities (vGlut2^+^/Homer1^+^) at both P10 and P21 in the hippocampal striatum lacunosum-moleculare layer (Fig. 5A-D). However, we observed no change in vGlut1^+^/Homer1^+^ synapse density at either P10 or P21 in hippocampus striatum radiatum layer (Fig. 5E-H). Taken together, our data demonstrate that microglial GPR56 also plays an important role in hippocampal synapse development.

**Fig. 5.**
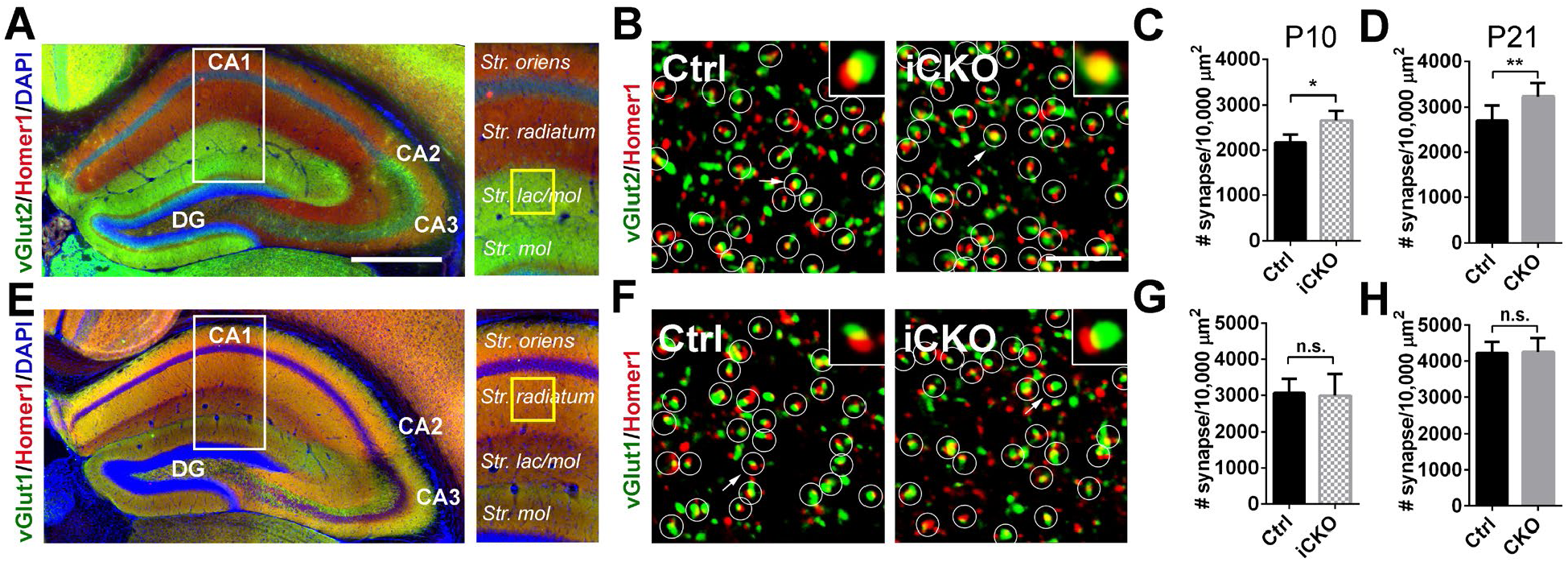
Microglial GPR56 is required for synaptic development in the hippocampus. (A) Representative images show hippocampal vGlut2 and Homer1 immunostaining. White box outlines the region of interest and yellow box shows the regions where confocal images were taken. Scale bar, 500 µm. (B) Confocal images of synaptic immunostaining in CA1 striatum lacunosum-moleculare (str. lac/mol) at P10. Arrows pointing to the enlarged synapse in the insets. Scale bar, 5 µm. (C, D) Quantification of synapse density in CA1 str. lac/mol in iCKO versus control at P10 (C), and CKO versus control at P21 (D). At P10, N(Ctrl) = 4, N(iCKO) = 3, p = 0.02. At P21, N = 8, p = 0.005. (E) Representative images show hippocampal vGlut1 and Homer1 immunostaining. (F) Confocal images of synaptic immunostaining in CA1 striatum radiatum at P10. (G, H) Quantification of synapse density in CA1 striatum radiatum in iCKO versus control at P10 (G), and CKO versus control at P21 (H). At P10, N = 5, p = 0.83. At P21, N = 5, p = 0.89. *p < 0.05, **p < 0.01, Student’s t-test. All data are presented as mean ± SD.

### Increased synaptic density correlates with functional consequences

During early postnatal periods, overlapping inputs from both eyes undergo a process of remodeling called eye-specific segregation, resulting in the termination of ipsilateral and contralateral inputs in distinct non-overlapping domains in the mature dLGN (Sretavan and Shatz, 1984). Defective synaptic pruning of RGC inputs to the relay neurons in the dLGN leads to incomplete eye-specific segregation (Chung et al., 2013; Schafer et al., 2012; Vainchtein et al., 2018). To investigate the effects of microglial GPR56 deficiency on eye-specific segregation, we performed anterograde tracing of RGC inputs with two different colored fluorescent CTB (Fig. 6A), and quantified the extent of segregation with a threshold-independent method as described (Tiriac et al., 2018; Torborg and Feller, 2004). We observed a significantly larger overlap at P10 in CKO mice (Fig. EV5A and B), persisting to P30 (Fig. 6B-D) when eye-specific segregation should be completed, suggesting improper organization of retinogeniculate projections sustained throughout development.

**Fig. 6.**
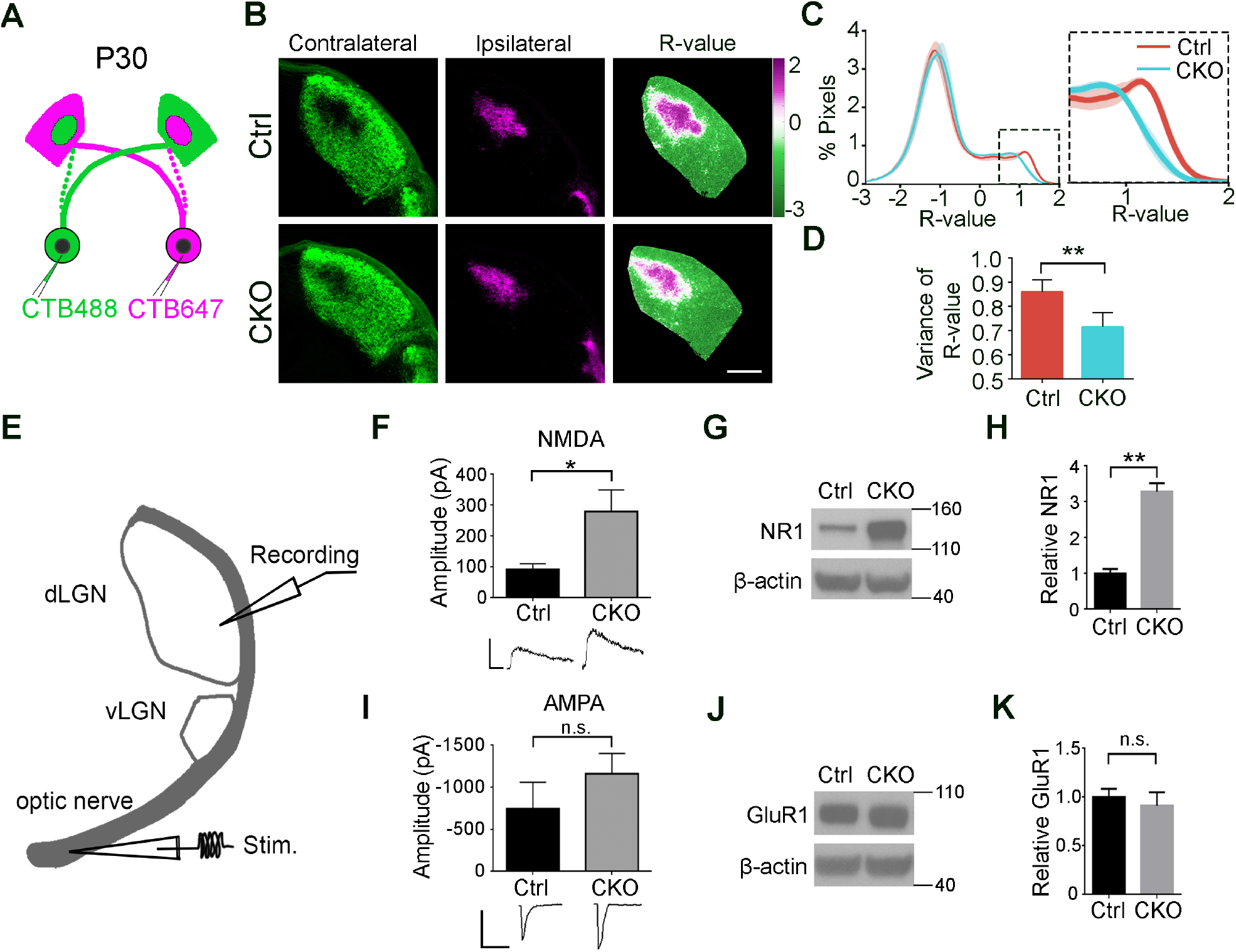
Microglial *Gpr56* deficiency leads to impaired retinogeniculate circuit organization and function. (A) A diagram of eye-specific segregation assay at P30. (B) CTB-labeled dLGN shows reduced eye-segregation at P30 in CKO mice. The left column shows contralateral dLGN labeled with CTB488 (green), and the middle one is ipsilateral dLGN with CTB647 (magenta). The right column represents the dLGN pseudocolored according to the R-value for each pixel (R = log(F_ipsi_/F_contra_)). Scale bar, 200 µm. (C) A histogram distribution chart of R-value for all pixels within dLGN represents the degree of eye-specific segregation. When R is 0, it means equal ipsilateral and contralateral fluorescence intensity at a pixel. A greater R-value means a bigger difference of ipsi-to-contraleteral fluorescence intensity. The narrower distribution of CKO in the inset indicates reduced segregation. (D) The variance of R distributions in P30 control and CKO mice. N = 4 (Ctrl), N = 6 (CKO), p = 0.004, mean ± SD. (E) A schematic diagram of electrophysiological recording in a parasagittal dLGN. (F) Maximal NMDAR mediated currents measured in the dLGN of P28∼P34 mice, p = 0.03, N = 14 (Ctrl), 23 (CKO) cells from 5, 7 mice, respectively. Mean ± SEM. (G) Western blot of NMDAR1 using microdissected dLGN tissue from P30 mice. (H) Quantification of NMDAR1 expression in CKO mice and control. N = 3, p = 0.007, mean ± SD. (I) Maximal AMPAR mediated currents measured in the dLGN of P28∼P34 mice. P=0.302, N = 13 (Ctrl), 17 (CKO) cells from 5, 7 mice, respectively. Mean ± SEM. (J) Western blot of GluR1 with microdissected P30 dLGN tissue. (K) Quantification of GluR1 expression in CKO mice and controls. N = 3, p = 0.40, mean ± SD. *p < 0.05, **p < 0.01, Student’s t-test.

To evaluate whether these excess synapses carry functional consequences, we examined the strength of the synaptic drive from RGCs to dLGN neurons by electrophysiological recordings. Whole-cell patch-clamp recordings from dLGN neurons were conducted in acute slices from either control or CKO mice at ∼P30 (Fig. 6E). When we examined maximal AMPA and NMDA receptor-mediated currents evoked by stimulation of the optic fibers that project to the dLGN, CKO mice displayed a significant increase in maximal NMDA receptor-mediated current (Fig. 6F), consistent with an increase in overall number of retinal inputs onto dLGN relay neurons (Fig. 4C-J) and increased NMDAR1 protein levels in western blot analyses of microdissected dLGN from P30 CKOs compared to those from controls (Fig. 6G and H). Mean maximal AMPA receptor-mediated currents were comparable between CKO and control mice (Fig. 6I), consistent with a comparable AMPAR protein level on western blot (Fig. 6J and K). Together, our data show microglial GPR56 is required for proper retinogeniculate circuit organization and function.

### Microglial GPR56 is required for the engulfment of PS^+^ synapses

Our data so far showed significant increases in selected synapses in CKO mice. To investigate whether this change results from decreased synaptic pruning by microglia, we performed *in vivo* engulfment assay. Compared to controls, the engulfment of RGC material in CKO microglia was decreased by 25.7% (Fig. EV5C and D). This change corresponded in magnitude to the increase in synapses at P5 (Fig. 4D) and supports the hypothesis that a reduction in the engulfment of RGC inputs by *Gpr56*-deficient microglia led to the overall increase in retinogeniculate synapses.

Considering that PS binds GPR56 and flags RGC presynaptic inputs for removal by microglia, we further hypothesized that the elevated synapse numbers may have resulted from a reduction in microglial engulfment of PS^+^ synapses. To test this hypothesis, we first examined the density of PS^+^ RGC inputs by PSVue labeling in the P6 dLGN of both CKO mice and controls. As predicted, we observed a significantly higher percentage of RGC inputs that were colocalized with PSVue in CKO, compared to controls (Fig. 7A and B). Considering the impaired ability to engulf synaptic elements by *Gpr56* CKO microglia (Fig. EV5C and D), the increase of PS^+^ RGC inputs in CKO dLGN might be a result of a decreased removal of PS^+^ RGC inputs by CKO microglia. To test this hypothesis, we performed the modified microglial engulfment assay in CKO mice and their controls. Indeed, we found the engulfment of PS^+^ RGC inputs was significantly decreased in CKO microglia, compared to controls (Fig. 7C and D). In contrast, the engulfment of PS^-^ RGC inputs was comparable between CKO and control microglia (Fig. 7E), Of note, *Gpr56* CKO microglia still contained a low level of PS^+^ RGC inputs, indicating that pathways other than GPR56 might mediate this process. Taken together, our data support the notion that GPR56 specifically regulates engulfment of PS^+^ synapses in microglia.

**Fig. 7.**
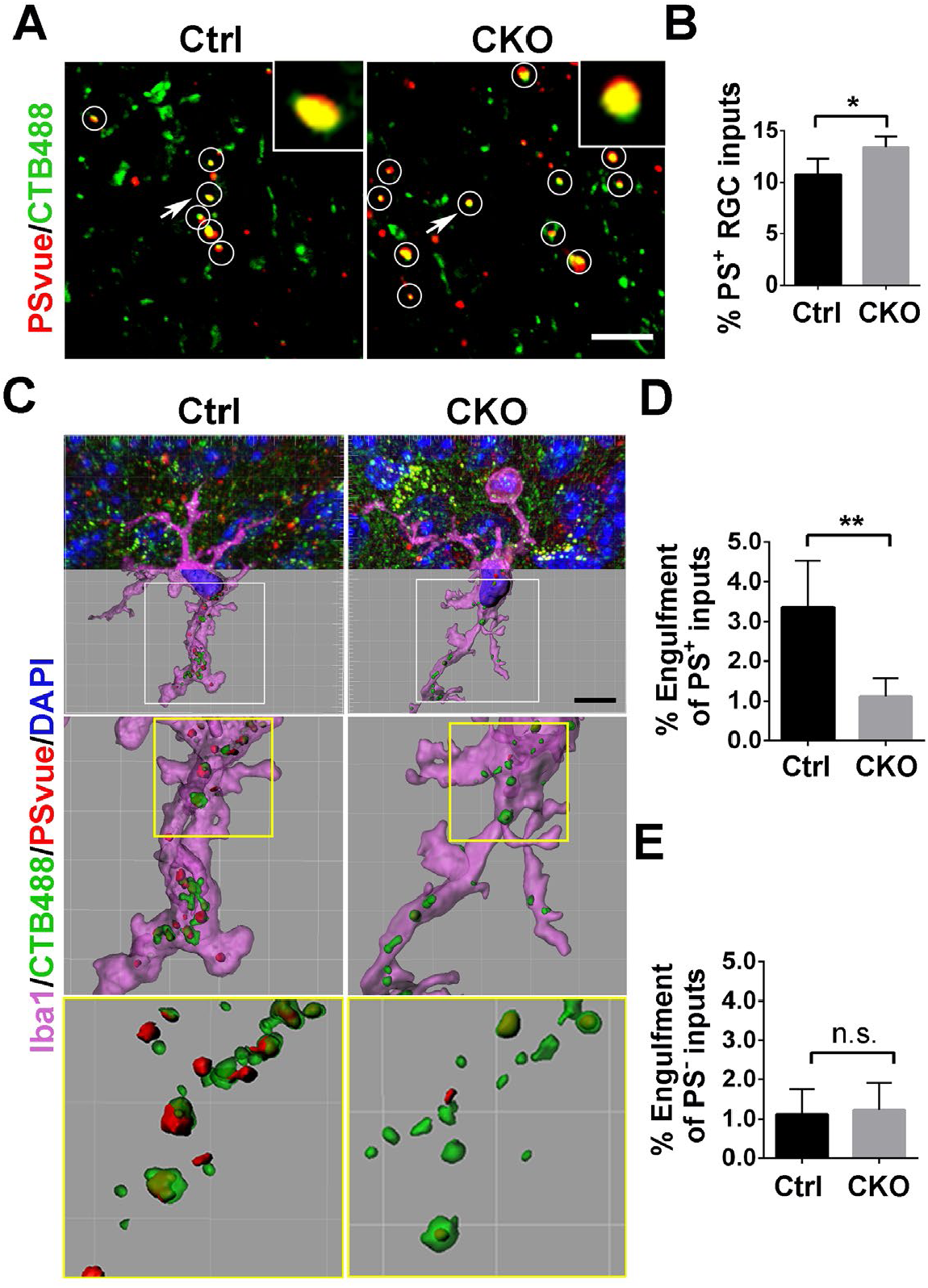
Deletion of microglial GPR56 causes decreased engulfment of PS^+^ RGC inputs. (A) Colocalized RGC inputs and PSVue signals in P6 control and CKO. White circles indicate PS^+^ RGC inputs. Arrows pointing to the RGC inputs enlarged in the insets. Scale bar, 5 µm. (B) The percentage of PS^+^ RGC inputs in total inputs in CKO and controls. N = 5, *p = 0.02. (C) Representative images of engulfed PS^+^ and PS^-^ RGC inputs by microglia in control and CKO. Scale bar, 10 µm. (D) Quantification of engulfed PS^+^ RGC inputs by microglia, which is calculated as: Volume of engulfed PS^+^ RGC inputs / Volume of microglia cell. (E) Quantification of engulfed PS^-^ RGC inputs in control and CKO. N=4 (Ctrl), N=5 (CKO), **p = 0.005 in (D), p = 0.80 in (E). * p < 0.05, **p < 0.01, Student’s t-test. Data are presented as mean ± SD.

## Discussion

### Alternative splicing expands the functional repertoire of GPR56 in brain development

Cells use alternative splicing to expand their regulatory and functional capacity of genomes (Braunschweig et al., 2013). GPR56 is a well-suited example in this regard. *Gpr56 S4* is an alternative splicing isoform that includes a 5’ deletion in exon 2, thus generating a new frameshifted transcript that uses a cryptic start site in exon 4 (Kim et al., 2010). As a consequence, the NTF of GPR56 S4 contains GAIN domain but lacks the PLL domain (Salzman et al., 2016), which mediates interactions with its two known extracellular ligands, collagen III and TG2 (Luo et al., 2012; Salzman et al., 2016; Yang et al., 2011). Here, we uncover a distinct function of the GPR56 S4 isoform in microglia where it binds PS to mediate synaptic pruning. This discovery reveals a critical biological function for the GAIN domain. Prior to our work, the GAIN domain had been largely viewed as an aGPCR signature structure that mediates autoproteolysis during protein maturation process.

### GPR56 regulates complex brain development in cell type-, isoform-, and ligand-specific manner

The adhesion GPCR (aGPCR) family, the second largest subfamily of GPCRs, consists of 33 members in the human genome (Hamann et al., 2015). Being a major class of matrix receptors, aGPCRs play essential functions in development by mediating cell-cell and cell-matrix interactions (Hamann et al., 2015). One of the most intensively studied aGPCRs associated with human developmental disorders is *GPR56*. Loss of function mutations in *GPR56* cause a devastating brain malformation called bilateral frontoparietal polymicrogyria (BFPP) (Piao et al., 2004). MRI analyses of BFPP brains show compound malformation including cortical lamination defect and abnormal CNS myelination (Piao et al., 2005; Piao et al., 2004).

Microglia enter the developing brain prior to neural tube closure, where they proliferate and propagate along with the sequential processes of neurogenesis, synaptogenesis and pruning, and myelination (Reemst et al., 2016). Recent literature indicates that microglia play a key role in each step of brain development (Hagemeyer et al., 2017; Miyamoto et al., 2016; Parkhurst et al., 2013; Sato, 2015; Schafer et al., 2012). Microglial *Gpr56* expression is governed by a super enhancer suggesting that it might be implicated in microglia core functions (Gosselin et al., 2014). This hypothesis is supported by the finding that *Gpr56* defines yolk sac-derived microglia (Bennett et al., 2018). In the present report, we provide insight into the functional significance of this expression pattern by showing that microglial GPR56 mediates synaptic pruning in multiple regions of the developing brain. Importantly, we show that an alternative splicing isoform, GPR56 S4, is responsible for microglia-mediated synaptic elimination by binding to PS, while the full length GPR56 is required for its functions in cortical development by interacting with its ligand collagen III, and CNS myelination by mediating a tripartite signaling with TG2 and laminin (Fig. 3 and 8) (Giera et al., 2018; Luo et al., 2011; Luo et al., 2012; Salzman et al., 2016).

**Fig. 8.**
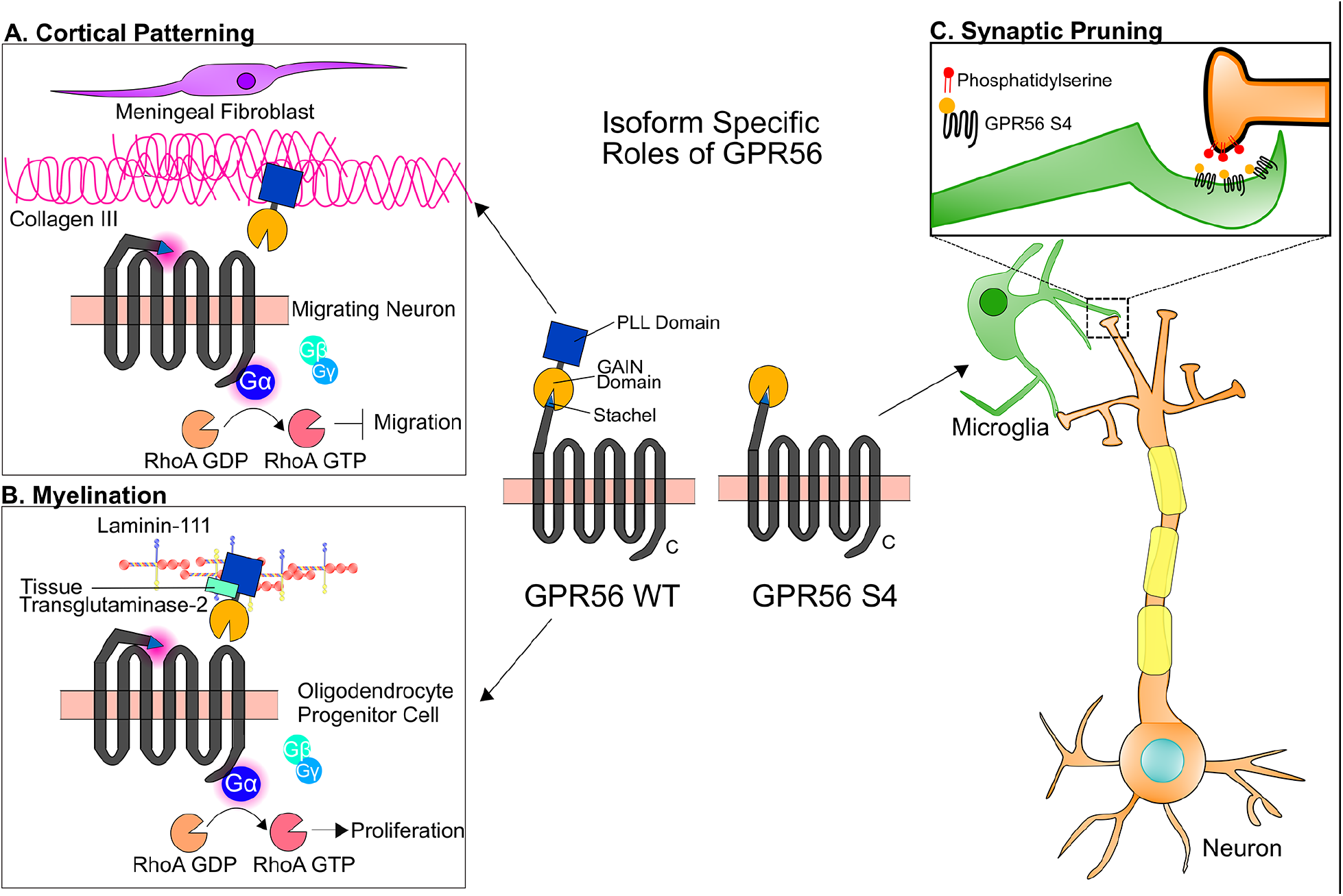
Isoform-specific roles of GPR56 in the brain. The NTF of WT GPR56 contains both PLL and GAIN domains. The S4 variant of GPR56 contains only the GAIN domain. (A) WT GPR56 participates in cortical patterning during brain development. Collagen III, secreted by meningeal fibroblasts, activates GPR56 by binding to its PLL domain and sends a stop signal to migrating neurons once they reached to the pial basement membrane (Luo et al., 2011). (B) In oligodendrocyte progenitor cells (OPCs), WT GPR56 binds to microglia-secreted tissue transglutaminase-2 (TG2) in the presence of laminin-111 to promote OPC proliferation (Giera et al., 2015; Giera et al., 2018). (C) In microglia, GPR56 S4 binds to externalized PS, which services as an “eat-me” signal for those synapses needed to be removed, and mediates the engulfment of synapses by microglia.

As previously reported (Giera et al., 2015), loss of GPR56 led to defects in myelination (Fig. 3G and H), which might impact synaptic pruning. However, in the developing visual system, myelination is initiated after P7 and doesn’t fully finish until P30 (Etxeberria et al., 2016; Mayoral et al., 2018). In contrast, synaptic pruning peaks at P5, and finishes around P10 in the dLGN. Here in *Gpr56 null* mice, we observed the defect of synaptic pruning as early as P5 (Fig. 3N and O), which occurred before myelination initialization. Based on these, it is unlikely that myelination defects impact synaptic pruning in this context.

### Exposed PS serves as an “eat-me” signal for synapses to be engulfed by microglia

Under normal conditions, PS is mainly restricted to the inner leaflet of the cell membrane. When cells undergo apoptosis, PS is exposed to the outer surface of the cell. This PS externalization is regulated by flippase and scramblase (Nagata et al., 2016; Zachowski et al., 1989). Caspases can inactivate flippase and activate scramblase at the plasma membrane by cleavage, and induce apoptotic PS externalization (Segawa and Nagata, 2015). Thus, exposed PS has become a classic “eat-me” signal for macrophage/microglia to phagocytose apoptotic cells.

It has been postulated that synapses could undergo focal apoptosis and be cleared. *In vitro* data showed that key characteristics of apoptosis, including PS externalization and caspase activation, were exhibited on cortical synaptosomes and on dendrites in cultured hippocampal neurons upon treatment of apoptosis inducer, like Staurosporine and Fe^2+^ (Mattson et al., 1998). Additionally, it has been shown that activated caspase-3 was colocalized with C1q on synapses in the brain tissue (Gyorffy et al., 2018) and that the focal activation of caspase-3 caused dendrite retraction and spine elimination without killing the neuron (Erturk et al., 2014). Thus, current literature supports that a focal apoptotic mechanism might underlie the synaptic pruning. However, there has been no report on the presence of PS^+^ synapses *in vivo*.

Here, for the first time, we demonstrate externalized PS on synapses *in vivo*, which strongly supports the focal synaptic apoptosis hypothesis. In the developing mouse visual system, we detected ∼10% of RGC synaptic inputs were PS^+^ at P6, an active period of microglia-mediated synaptic pruning; and much fewer PS^+^ inputs were found at P13, when synaptic pruning was essentially completed in the dLGN (Fig. 1C and D). This finding indicates that synapses intended to be removed expose PS as an “eat me” signal to enable a targeted removal by microglia. We further strengthened this notion by revealing that over 70% RGC synaptic inputs engulfed by microglia were PS^+^, in contrasted to 27% of PS^-^ RGC inputs (Fig. 1K). Taken together, our data support that exposed PS serves as an “eat-me” signal for synapses to be engulfed by microglia.

### GPR56 functions as a PS receptor

We uncover a new direct PS binding protein, the S4 isoform of GPR56. Several different PS recognizing membrane receptors have been identified under the categories of either direct PS receptors and indirect opsonin-based PS receptors (Bevers and Williamson, 2016). For example, BAI1 (Park et al., 2007), Tim-4 (Miyanishi et al., 2007), Stabilin-2 (Park et al., 2008) are direct PS receptor, whereas TAM receptors (Tyro3, Axl, and MerTK) bind PS via GAS6 and Protein S (Lemke, 2013; Mark et al., 1996; Nagata et al., 1996; Ohashi et al., 1995; Stitt et al., 1995). Most of these receptors are related to engulfment of apoptotic cells. One intriguing question is whether these PS receptors also mediate engulfment of synapses. Indeed, our present study demonstrates that PS binding protein GPR56 promotes the engulfment of PS^+^ synapses by microglia. Interestingly, we found CKO microglia still engulf some PS^+^ synapses (Fig. 7C and D), indicating there might be other molecular pathways that regulate engulfment of the remaining PS^+^ synapses.

Another provocative question is how GPR56 functions together with other PS receptors in regulating developmental synaptic pruning. One possible scenario is that various PS receptors function in concert to prune synapses, as observed in microglia engulfment of dying neurons where BAI1 regulates the formation of phagosomes around dying neurons and Tim-4 controls phagosome stabilization (Mazaheri et al., 2014). Another possibility is that different cell types use different PS receptors. In supporting the second possible scenario, single cell RNAseq database reveal that BAI1 and MEGF10 are highly expressed in astrocytes, but barely detectable in microglia, GPR56 and MERTK are expressed in both microglia and astrocyte, whereas Tim-4 and Stabilin2 are minimally detectable in microglia and astrocyte (Zhang et al., 2014).

### Concluding remarks

Defective synapse pruning has been implicated in autism spectrum disorder (Tang et al., 2014), whereas excessive synapse removal has been linked to schizophrenia (Sekar et al., 2016; Sellgren et al., 2019). Insight into the mechanisms underlying synapse refinement during development will help decipher neurodevelopmental disorders linked to synapse imbalance. Key unanswered questions include how synapses destined for removal externalize PS and whether GPR56 functions independently or in concert with complement components in microglia-mediated synaptic pruning. Given its high expression in adult microglia (Matcovitch-Natan et al., 2016), our data also raise a broader question of whether microglial GPR56 has any role in adult synapse homeostasis. Given its multifarious and cell-type specific roles in neurodevelopment (Folts et al., 2019), further study of GPR56 function in synaptic pruning represents an attractive opportunity to integrate this process into other aspects of brain wiring including myelination. This understanding will promote the objective of enabling therapeutic targeting of GPR56 for combating neurodevelopmental and neurodegenerative disorders.

## Materials and Methods

### Animals

All mice were handled according to the guidelines of Animal Care and Use Committee at Boston Children’s Hospital and University of California, San Francisco. *Gpr56*^*fl/fl*^ mice were generated as previously described (Giera et al., 2015). The *Cx3Cr1-cre* (B6J.B6N(Cg)-*Cx3cr1*^*tm1.1(cre)Jung*^/J, #025524) and *Cx3Cr1-creER* (B6.129P2(Cg)-*Cx3cr1*^*tm2.1(cre/ERT2)Litt*^/WganJ, #021160) mice were obtained from Jackson Laboratories. Considering both *Cx3Cr1*^*Cre*^ and *Cx3Cr1*^*CreER*^ are knock-in mice, we crossed these mice with *Gpr56*^*fl/fl*^ to generate *Gpr56*^*fl/fl*^;*Cx3Cr1-cre (or creER)*^*+/-*^ as conditional knockout mice, and *Gpr56*^*+/+*^; *Cx3Cr1-cre (or creER)*^*+/-*^ as controls. CreER is a tamoxifen-inducible Cre recombinase. To induce a deletion of microglial *Gpr56*, 40 µg tamoxifen (in corn oil, Sigma) per day for 3 consecutive days (P1-P3) were given to neonatal animals via intraperitoneal injection (Parkhurst et al., 2013). To generate *Gpr56 null* mice, *Gpr56*^*fl/fl*^ mice were crossed with *CMV-cre* mice (JAX stock #006054) (Schwenk et al., 1995) to delete exons 4-6, causing a deletion of G*pr56* in all tissues. Rosa^GFP^ reporter mice were made from pR26 CAG/GFP vector (Plasmid #74285, Addgene), which contains a loxP-flanked STOP cassette and a GFP reporter, which is expressed under the control of an IRES. Then the vector was micro-injected into C57BL/6 zygotes to make transgenic mice via CRISPR/Cas9 strategy in the Transgenic Core Laboratory of Boston Children’s Hospital (Chu et al., 2016).

### Immunohistochemistry

Mouse brains were collected following PBS perfusion and fixation with 4% PFA, and cryoprotected in 30% sucrose. OCT-embedded tissues were cryosectioned at 14 µm or 40 µm. For immunostaining, 14-µm or 40-µm Sections were incubated with blocking buffer (10% Goat serum + 1%BSA; 0.3% TritonX/PBS) for 2 hours and stained with primary antibodies overnight at 4°C (guinea pig anti-vGlut2, 1:1000, Millipore AB2251-I; guinea pig anti-vGlut1, 1:1000, Millipore AB5905; rabbit anti-Homer1, 1:250, Synaptic Systems, 160 003; rabbit anti-Iba1, 1:250, Wako 019-19741; guinea pig anti-Iba1, 1:500, Synaptic Systems 234 004; rat anti-CD68, 1:250, AbD Serotec MCA1957; rat anti-MBP, 1:100, Abcam ab7349), followed by fluorophore-conjugated secondary antibodies for 2 hours at room temperature. Tissues were mounted with Fluoromount-G (Southern Biotech, 0100-01) or Vectashield antifade mounting medium (Vector Laboratories, H-1000).

### In vivo phosphatidylserine labeling

P5 mice were given an intraocular injection of CTB647 into both eyes. Twenty-four hours later, mice were mounted on a neonatal mice adaptor (RWD Life Science, #68072), and given an intracranial injection of various labeling probes. The stereotaxic setting was −1.5 mm anteroposterior, 1.09 mm mediolateral and −2.15 mm dorsoventral to lambda. 2µl PSVue (20µM in TES buffer, Millipore Sigma T1375) or pSIVA(1:2, 2mM Ca^2+^ in HBSS) were injected into the gap area between hippocampus and dLGN via a 33G needle attached Hamilton syringe at 0.33µl/min (see Fig. EV1A and C). Six hours later, brains were collected freshly, and sectioned at 180µm in ice-cold TES buffer or 1mM Ca^2+^ HBSS on the vibratome (Leica VT1000 S). Two to three sections with medial dLGN were chosen, and directly mounted on homemade framed (concave, 200µm depth) glass slides. The images were taken immediately within an hour on a Zeiss LSM700 confocal microscope at 63X.

### *In vivo* engulfment

*In vivo* engulfment assay was carried out as previously described (Schafer et al., 2012). Briefly, P4 mice were given an intraocular injection of 0.5µl 0.5% CTB594 into each eye 24 hours prior brain collection. 40µm cryosections were obtained for the assay. 2-3 sections of medial dLGN were chosen from each mouse for further free-floating immunostaining with Iba1 antibody. Images were taken on a UltraView Vox spinning disk confocal microscope at 60X with 0.2µm z-step or on a Zeiss LSM700 confocal microscope at 63X with 0.2 µm z-step. ImageJ (NIH) and Imaris (Bitplane) were used for image processing. 3D reconstruction of microglia and RGC inputs were created by surface rendering in Imaris. To enable quantitative assessment of engulfment, and to control for variation in microglia volume, we calculated the engulfment percentage as: Volume of internalized RGC inputs / Volume of microglia. To quantify the engulfment of PS^+^ or PS^-^ RGC inputs, the following formula was used: % Engulfment of PS^+^ or PS^-^ RGC inputs = Volume of internalized PS^+^ or PS^-^ inputs / Volume of microglia.

### Generation of GPR56 fusion proteins

Human IgG Fc tagged constructs were generated by PCR. For GPR56 NTF-hFc, the PCR forward primer is: 5’-CCATGGAAGACTTCCGCTTCTGTGGCC-3’, the reverse primer is: 5’-CAGATCTCAGGTGGTTGCACAGGCAGG-3’. For GPR56 GAIN-hFc, the PCR forward primer is: 5’-ATCCATGGTTATGTG TGATCTCAAGAAGGAATTGC-3’, the reverse primer is: 5’-TCTAGATCTCAGGTGGTTGCACAGGCAGG-3’. Then PCR product was inserted into pFUSE-hIgG2-Fc2 (InvivoGen, Cat. pfuse-hfc2) vector between Nco1 and BglII sites. For the expression of GPR56 fusion proteins, the above constructs were transiently transfected into HEK-293T cells (ATCC). 24 hours later, the culture media was changed to serum-reduced Opti-MEM (Gibco, 31985070). During incubation, GPR56 fusion proteins would be secreted into the culture media. This conditioned media containing fusion proteins was harvested 48-72 hours later, and concentrated as previously described (Li et al., 2008). The proteins were purified using HiTrap protein A column (GE Healthcare, 17040303).

### Flow cytometry

GPR56 NTF-hFc, GAIN domain-hFc, or hFc were labeled with Alexa Fluor 647 (AF647) using the Alexa Fluor 647 NHS Ester labeling kit (ThermoFisher Scientific, Cat #A20006) according to the manufacturer’s protocol. Briefly, 75 µM protein was incubated with 300 µM AF647 at room temperature for 4 hours, followed by 200 mM Tris to quench the reaction. The whole reaction mixture was run through a Sephadex G-50 column and protein-dye conjugate was collected. Dye labeling efficiency for each protein ranged from 1.44-1.46 moles of dye per mole of protein.

Ba/F3 cell line was cultured and passaged as described (Cermak et al., 2016).It was maintained in RPMI-1640 media with L-Glutamine (Invitrogen, Cat #11875093) supplemented with 10% FBS, Penicillin-Streptomycin, and kept at 37°C, 5% CO2 in a humidified incubator. Cells were harvested and washed twice with ice-cold Hank’s Balanced Salt Solution (HBSS) and resuspended in HBSS. Cells were treated with 1 µM calcium ionophore A23187 (Millipore Sigma, Cat #C7522) at 37°C for 15 minutes, with gentle agitation every 5 minutes, followed by washing with HBSS. To measure direct binding, cells were resuspended in 0.9% NaCl and incubated with 1 µM protein-Alexa Fluor 647 conjugate at room temperature for 2 hours. After binding, cells were washed once and resuspended in HBSS, followed by flow cytometry on an LSRFortessa (BD Biosciences). For the competition assay, cells were treated with A23187 as above and washed with HBSS. Next, cells were resuspended in 100 µL staining buffer (10 mM HEPES, 140 mM NaCl, 2.5 mM CaCl_2_, pH 7.4) and incubated with 1 µM AF647 conjugated NTF, GAIN, or hFc and 50 nM Annexin V-FITC for 1 hour at room temperature. Cells were washed once with HBSS, followed by flow cytometry on an LSRFortessa (BD Biosciences). Ba/F3 cell line was a gift from the Manalis laboratory at Massachusetts Institute of Technology.

### Protein lipid overlay assay

The membrane lipid strips (Echelon Biosciences, Inc.) were blocked in 3% fatty-acid-free BSA and 1% fatty-free milk in 0.1% Tween-20 TBS for 1 h at room temperature. The membrane was then incubated in 3%BSA containing 3 μg/ml purified human Fc-tagged GPR56-GAIN domain (hFc-GAIN) at 4 °C with gentle agitation. After three times wash, the strips were incubated with goat anti-human IgG Fc HRP-conjugated secondary antibody for 1h (Thermo Fisher Scientific). The film was exposed following routine immunoblotting steps. The whole experiment was processed in the dark, because this lipid strips were sensitive to light.

### QPCR

RNA of microglia was isolated using the miRCURY™ RNA isolation kit (EXIQON, Cat. #300110), and was reverse-transcribed using a kit of SuperScript™ IV VILO™ Master Mix with ezDNAse (Invitrogen, Cat. # 11766050). CDNA was used as template for PCR reactions with LightCycler® 480 SYBR Green I Master (Roche, Cat. # 04707516001) in a Roche LightCycler® 480 II machine. Forward and reverse primers for qPCR1 (338-421) are 5’- GCGGCAGATGGTCTACTTCCT-3’ and 5’-AAGCGGAAGTCTTCTCGGGG-3’, respectively. Forward and reverse primers for qPCR2 (1585-1669) are 5’- TTGCTGCCTACCTCTGCTCC-3’ and 5’- AGCAGGAAGACAGCGGACAG-3’, respectively.

### Synapse quantification

Confocal microscopy images were obtained with Zeiss LSM 700 System. For synapse quantification in dLGN, medial dLGN slices were used. In each dLGN, three fields of view (5 serial optical sections, 0.5 µm Z-step, 101.5 µm * 101.5 µm, 1024 * 1024 pixel) were acquired in the upper part of the core region using a 63X/1.40 oil objective. Colocalization of vGlut2/Homer1 or vGlut1/Homer1 was quantified as described (Lehrman et al., 2018). The whole process was run in the ImageJ software (NIH, Bethesda, MD). First, each channel’s background was subtracted with a rolling ball radius of 10 pixels. Then thresholding was properly applied to each channel to distinguish synaptic puncta from background, and generate two new binary images. To detect the overlay of pre- and post-synaptic puncta, a logical operation “AND” was performed between these two images. Lastly, the overlaid puncta were counted as synapse using the “analyze particles” function. The full code can be found at: https://github.com/TaoLi322/Microglia_GPR56_Synapse/blob/master/Synapse-quantification. All images were acquired and analyzed blindly.

For super-resolution imaging of synapse s, a modified version of the protocol in Hong et al. 2017 (Hong et al., 2017) was employed. In brief, imaging was performed using a Zeiss ELYRA PS1 structured illumination microscopy (SIM). Tissue were mounted with Prolong Gold (Invitrogen P36934) and covered by High Precision Cover Glass (1.5H, Azer scientific). Sections were imaged at 90 nm Z-step using a 100X oil-immersed objective lens. Images were processed using the Zen image software (Carl Zeiss), and then inputted into Imaris for 3D rendering and analysis. The vGlut2 channel was processed using surface rendering function, and Homer1 channel was processed with the spot function. A MatLab program “spot to surface” was applied to count the number of adjacent Homer1 spots to vGlut2 surface (≤ 100 nm, 200 nm and 300 nm distance from spot centers to surface).

### RNAscope

RNAscope was performed using RNAscope® Multiplex Fluorescent Reagent Kit v2 for fixed and frozen 12um thick sections according to the manufacturer’s instructions. RNAscope^®^ Probe-Mm-*Gpr56* (Cat No. 318241) was used to detect expression of the C-terminal target region of *Gpr56* followed by immunohistochemistry for Iba1 (1:400, Wako, 019-19741). In short, after signal amplification step for *Gpr56*, sections were permealized using 0.3% Triton-X 100 in PBS for 10 mins, followed by blocking with 10% goat serum, 1% BSA and 0.1% Triton-X 100 in PBS for 1 hr at RT and incubating the primary antibody IbaI in the blocking buffer overnight at 4°C. Appropriate secondary antibody was used to visualize Iba1 expression.

### Western blot of dLGN

Mice (P30) were anesthetized and decapitated. Mouse brains were immediately removed into ice-cold HBSS and coronally sectioned at 200μm using vibratome. DLGN were dissected out under dissection microscope, and homogenized in proteinase inhibitor-added RIPA buffer. After 30min incubation on ice, samples were centrifuged at 14,000xg for 10min. Supernatant were collected and used for WB of NMDAR1 (anti-NR1, 1:2000, Sigma-Aldrich 05-432) and vGlut2 (anti-vGlut2, 1:5000, Millipore AB2251-I). For WB of GluR1 (anti-GluR1, 1:1000, Cell signaling 13185S), supernatant was boiled for 5min before loading.

### Microglia cell isolation

Microglia were isolated from whole brains without cerebellum using a modified Percoll gradient assay (Frank et al., 2006). The whole process was carried out under ice-cold conditions, unless other temperatures were used. Briefly, anesthetized mice were perfused with ice cold Hank’s balance salt solution (HBSS). The fresh brains were immediately minced into 1-2 mm chunks using a clean razor blade, and transferred to a Dounce homogenizer. Using a loose glass pestle, brain tissues were gently homogenized with around 15 strikes, until large chunks had dissociated. Then repeat the homogenization with a tight pestle until the tissue had been mostly dissociated to single cells. The cell suspension was pelleted at 300xg for 7 min. The cell pellets were resuspended in 5 ml of 70% Percoll (GE healthcare, Cat. # 17089101) diluted in HBSS. Another layer of 5 ml 37% Percoll was gently added on top of the cell suspension layer. Then it was spun at 800xg, 22°C for 25 min, with the acceleration rate setting at 5 and deceleration rate at 1. A thin, cloudy layer of cells at the interface was carefully collected and pelleted. These cells were used for following western blot of microglia.

The cells were resuspended in FACS buffer (0.2% BSA in HBSS), and stained with CD11b-PE (BD, Cat. #553311) and CD45-BV421 (BD Horizon™, Cat. #563890) at a 1:100 dilution for 20min. The cells were then washed and spun down at 300xg for 7min. The samples were resuspended again in FACS buffer containing DRAQ7 (Abcam, cat. #ab109202) at a 1:100 dilution, and filtered through a 35 µm cell strainer. Using a BD FACSAria II sorter, CD11b^+^, CD45^low^, DRAQ7^-^ cells were collected as microglia, which were further used for qPCR experiments.

### Eye specific segregation

Threshold-independent analysis of eye-specific segregation was performed as described before (Torborg and Feller, 2004). Mice were anesthetized with isoflurane during the whole procedure. 3ul 0.2% cholera toxin B subunit (CTB) conjugated Alexa 488 dye (CTB488, Life Technologies, C22841) was intravitreally injected into the left eye, and CTB conjugated Alexa 647 dye (CTB647, Life Technologies, C34778) into the right eye of P30 mice. 24 hours later, mice were transcardially perfused with PBS and fixed with 4% PFA, and cryoprotected in 30% sucrose. The injection/labeling efficiency was checked by visualizing retinas and superior colliculus (Appendix Fig. S8A). Brains were cryosectioned coronally at 60 μm and mounted with Fluoromount-G (SouthernBiotech). Since variance of R-value varies much from cranial to caudal LGN (Tiriac et al., 2018), we chose three to four dLGN (420 to 600 µm from caudal LGN) as indicated in (Appendix Fig. S8B) for analysis. 16-bit images were digitally acquired at 20X with a CCD camera (Nikon eclipse Ti). The same exposure time and gains were used for each channel across all samples. For P30 mice, images were analyzed in a threshold-independent way. First, background fluorescence values were calculated from non-LGN regions, and used to be subtracted from images. Then images were normalized by adjusting the contrast to 5% saturation. For each pixel, R-value was computed as R = log_10_(F_ipsi_ / F_contra_), where F_ipsi_ or F_contra_ is the fluorescence intensity of the pixel in ipsilateral or contralateral channel, respectively. Those pixels whose values equal to zero were excluded. Over 99.5% of R-values ranged between −3 to 2, and they were plotted as a distribution graph. The variance of R-values was computed to quantify the extent of segregation. The greater variance means more segregation between ipsilateral and contralateral RGC inputs. The full code can be found here: https://github.com/TaoLi322/Microglia_GPR56_Synapse/blob/master/Eye-Seg_R-value.

For P10 mice, 1µl of 0.2% CTB488 and CTB594 (Life Technologies, C22842) was injected into the left or right eye, respectively. The percentage of overlapped left and right eye projections in dLGN was quantified using a multi-threshold quantitative method, as described before (Torborg and Feller, 2004).

### LGN slice preparation and electrophysiology

Mice (P28–P34) were decapitated; their brains were removed and placed in a 4°C choline solution, containing: 78.3 mM NaCl, 23 mM NaHCO_3_, 23 mM glucose, 33.8 mM choline chloride, 2.3 mM KCl, 1.1 mM NaH_2_PO_3_, 6.4 mM MgCl_2_, and 0.45 mM CaCl_2_. The two hemispheres were then separated by an angled cut (3°–5°) relative to the cerebral longitudinal fissure. The medial aspect of the right hemisphere was then glued onto an angled (20°) agar block, and 250 μm LGN slices were cut in ice cold, choline solutions using a vibratome (Leica VT1200S) based upon previously described protocols (Chen and Regehr, 2000; Turner and Salt, 1998). Slices were allowed to recover at 33° C for 30 minutes in choline solution and then for an additional 30-50 minutes in artificial cerebral spinal fluid (ACSF). ACSF contained: 125 mM NaCl, 25 mM NaHCO_3_, 25 mM glucose, 2.5 mM KCl, 1.25 mM NaH_2_PO_3_, 1 mM MgCl_2_, and 2 mM CaCl_2_. All solutions were continuously supplied with oxygen (95% O_2_/5% CO_2_).

Whole-cell voltage-clamp recordings of neurons from the dLGN were conducted using a Multiclamp 700B amplifier, digitized with a Digidata 1440A, collected with Clampex 10.2, and analyzed using Clampfit 10.2 (Axon Instruments). The slice containing dLGN was perfused with oxygenated ACSF, supplemented with 50 µM picrotoxin. Neurons were then patched using glass electrodes (3–4 MΩ) filled with 35 mM CsF, 100 mM CsCl, 10 mM EGTA, 10 mM HEPES, and 0.1 mM D600 (methoxyverapamil hydrochloride). Internal solution has a pH of 7.32 with CsOH. ACSF filled glass pipettes were placed in the optic tract, and these stimulating electrodes were moved repeatedly until reaching the location that gave the largest postsynaptic response. The maximum AMPA and NMDA currents were then determined by gradually increasing the stimulus intensity from 0.1 up to 1 mA. The maximum response size was considered to be the amplitude of the response after 3 consecutive increases in stimulation intensity failed to result in a larger response, or if the response amplitude decreased with an increase in stimulation intensity. AMPA responses were collected at −70 mV and NMDA responses at +40 mV, and were collected with alternate stimulations. To confirm that the optic tracts and not cortical inputs were stimulated, paired pulse data with a 50 ms interstimulus interval were collected, given that optic inputs usually demonstrate paired pulse depression, and cortical inputs paired pulse facilitation. No difference in paired pulse depression was seen between control and conditional knockout mice (Appendix Fig. S9, p = 0.694, Student’s t-test).

### Statistical Analysis

For all quantification, images were acquired blindly to genotype before quantification. All data are shown as mean ± SD or mean ± SEM as indicated in figure legends. Asterisks indicate significance: ****p < 0.0001, ***p < 0.001, **p < 0.01, *p < 0.05. All effects of genotype were analyzed by Student’s t-test, one-way ANOVA, or two-way ANOVA (Graph Pad Software, Inc).

## Data and materials availability

None.

## Acknowledgments

We thank Dr. Chinfei Chen at Boston Children’s Hospital for technical support on electrophysiology and thoughtful discussion on the manuscript, Douglas Richardson at Harvard Imaging Core for kind technical support on SIM imaging, Alexandre Tiriac and Marla Feller for kindly providing technical support on eye segregation analysis, Aashish Manglik for the Alexa Fluor 647 NHS Ester labeling kit and technical support, Dr. Scott Manalis for the Ba/F3 cell line, Dr. Greg Lemke at Salk Institute for sharing GAS6 plasmids, Nicole Scott-Hewitt for thoughtful discussion, and Dario Tejera, Eric Huang, Anna Molofsky, Mercedes Paredes, and Kelly Monk for critical reading of the manuscript.

## Author contributions

T.L. and X.P. conceived and designed the experiments, performed data analyses, and wrote the manuscript. T.L. Performed most experiments. B.C., C.G., T. K., S.G., R.L., D. Y. and H. O. contributed to experiments and data analysis. E.J.-V. and H.U. designed, performed, and analyzed the electrophysiology experiments. A.M. and B.S. assisted on synapse quantification and *in vivo* engulfment assay. All authors read and edited the manuscript.

## Supplemental data

### Expanded View Figures

**Fig. EV1.**
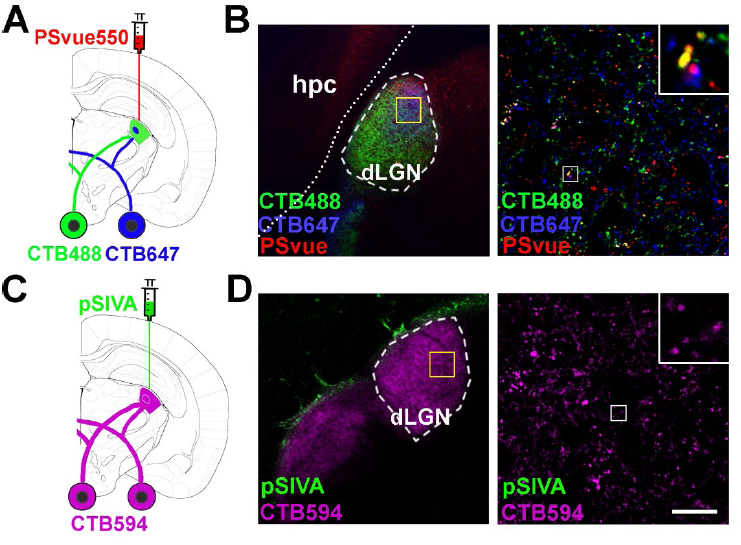
RGC synaptic inputs were labeled by PSVue. (A) A diagram illustrating PS labeling by PSvue550 and RGC inputs anterograde tracing by CTB. CTB was intraoccularly injected 24 hours prior to PSVue/pSIVA injection. (B) Left panel shows well-diffused PSVue into dLGN. The yellow box indicates the region where the images were taken. Right panel shows RGC inputs colocalize with PSVue signal. The white box indicates the region of higher magnification image shown. (C) A diagram showing pSIVA labeling and RGC inputs tracing by CTB. (D) Left panel shows pSIVA accumulated in the border between hippocampus and LGN. Right panel shows minimal pSIVA colocalized with RGC inputs. Scale bar, 20 µm.

**Fig. EV2.**
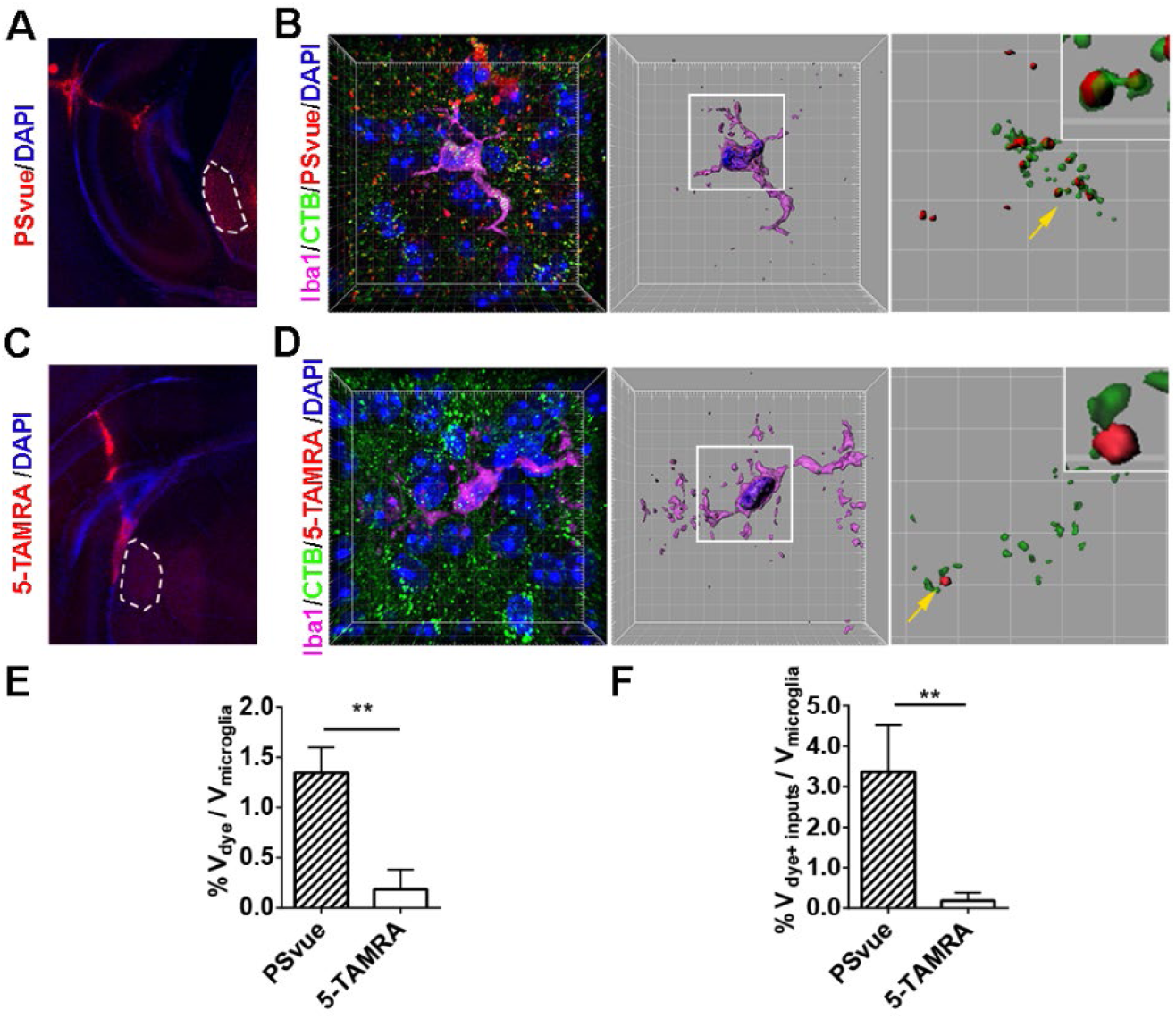
Microglia engulf PSVue-labeled RGC inputs. (A) An image shows the PSVue signal after intracranial injection. (B) A representative image of microglia from PSVue treated dLGN. Nuclei were labeled with DAPI. The mid panel shows a 3D surface rendered microglia (purple) with DAPI (blue) and engulfed inputs (green) and PSVue (red). The right panel shows engulfed RGC inputs and PSVue inside of microglia.The magnified insert shows PS^+^ RGC inputs. (C) An image shows the 5-TAMRA signal after injection. (D) A representative image of microglia from 5-TAMRA treated dLGN. Very few 5-TAMRA puncta were observed. The middle panel shows a surface rendered microglia with engulfed inputs (green) and 5-TAMRA (red). The right panel shows engulfed RGC inputs and 5-TAMRA. The insert shows RGC inputs do not overlap with 5-TAMRA. (E) Quantification of engulfed PSVue and 5-TAMRA dyes by microglia. N=4 (PSVue), N=3 (5-TAMRA), **p = 0.001. (F) Quantification of engulfed PSVue-positive or 5-TAMRA-positve RGC inputs by microglia. N=4 (PSVue), N=3 (5-TAMRA), **p = 0.006 by Student’s t-test. Data are presented as mean ± SD.

**Fig. EV3.**
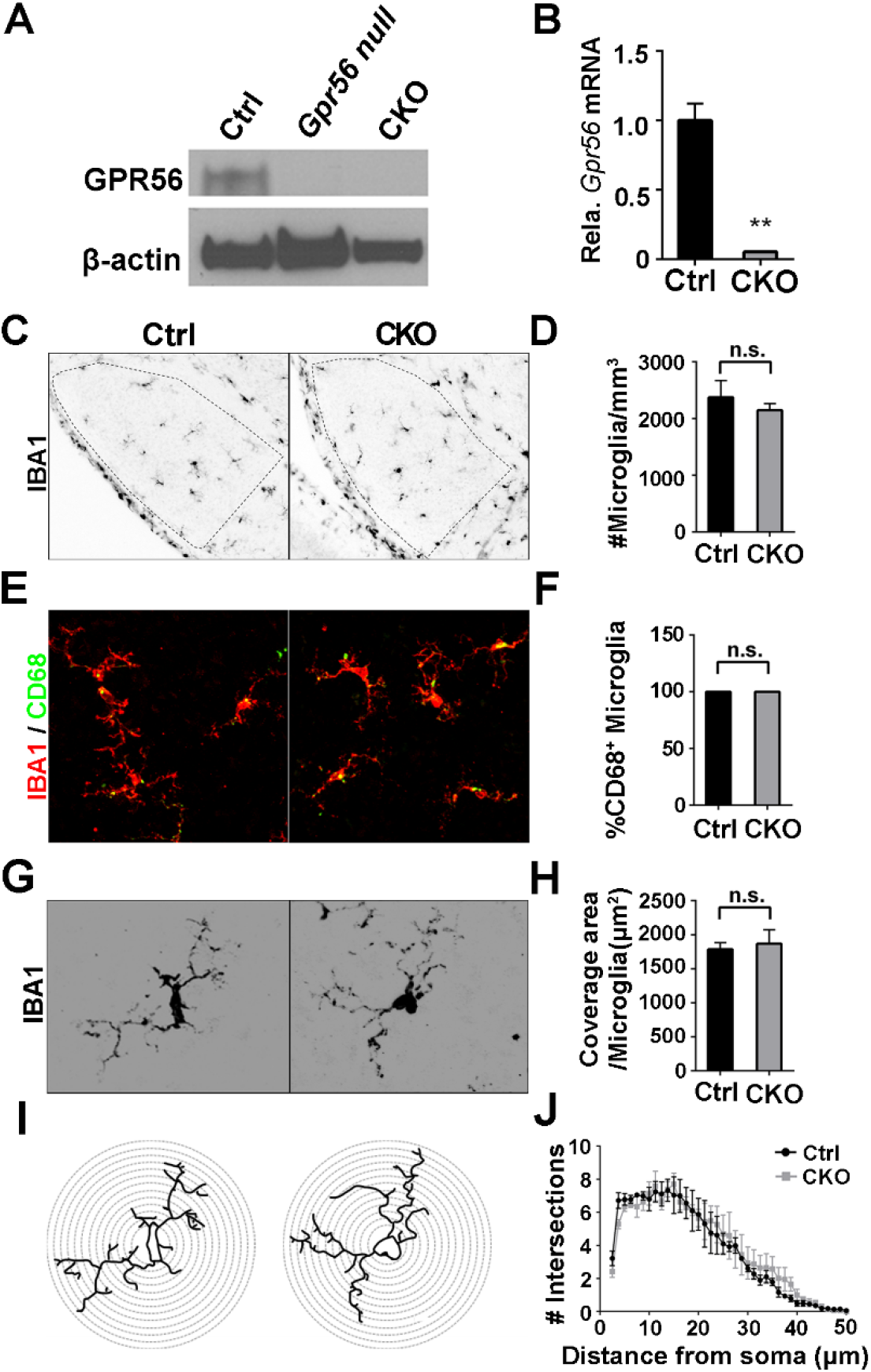
No significant change of cellular properties after deleting microglial *Gpr56*. (A) Western blot shows GPR56 protein was detected in the isolated microglia from control mice, but not in the microglia from *Gpr56 null* or CKO mice. (B) QPCR detects a high level of *Gpr56* transcripts in control microglia, compared to CKO microglia. N=3, **p < 0.01 by Student’s t-test. Data are presented as mean ± SEM. (C) Images of microglia stained by anti-Iba1 in dLGN at P5. (D) Quantification of microglial density between CKO and controls. N = 3 (Ctrl), N = 4 (CKO), p = 0.462. (E) Representative images of Iba1 and CD68 double staining. (F) Quantification of the percentage of CD68 positive microglia. 100% microglia are CD68 positive in both CKO and controls. N = 3. (G) Images of individual microglia, using Iba1 immunostaining to visualize its morphology. (H) Quantification of coverage area of each microglia between CKO and controls. N = 35 (Ctrl), N = 22 (CKO) microglia from 3, 4 mice, respectively. p = 0.685, Student’s t-test. (I) Images depicting concentric circles upon manually outlined microglia at 1.25 μm intervals for Sholl analysis. (J) Sholl analysis shows no significant change in arbor complexity in CKO. N = 3, F (1, 156) = 0.88, P = 0.35 by two-way ANOVA with Bonferroni’s post hoc test. Data are presented as mean ± SEM.

**Fig. EV4.**
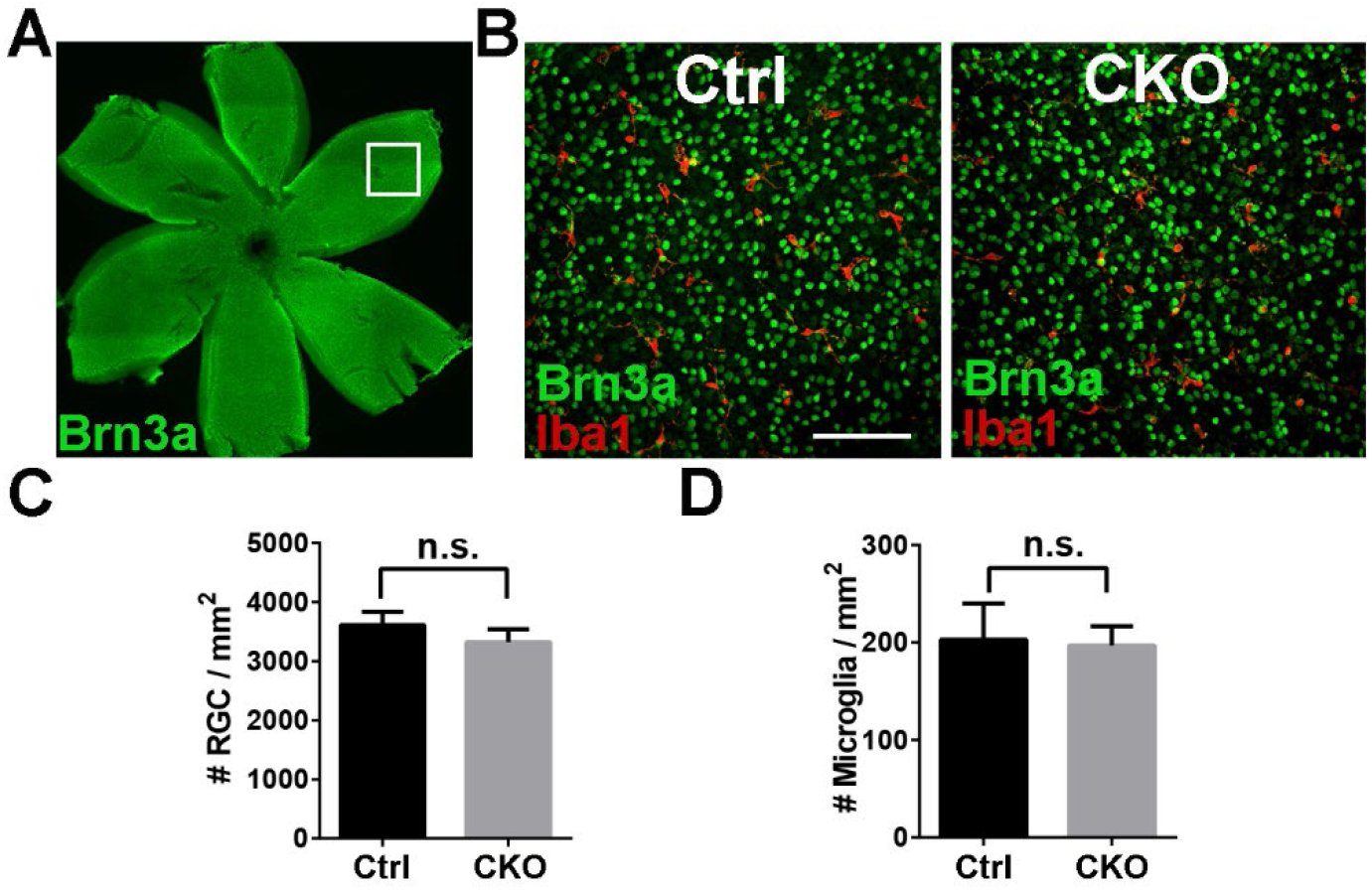
Deleting microglial *Gpr56* has no significant effect on RGC density in P5 retina. (A) Whole mount retinal staining of RGC using Brn3a antibody. (B) Representative images of RGC and microglia staining using Brn3a and Iba1 antibodies in retina. (C) Quantification of RGC density in controls and CKO. (D) Quantification of retinal microglia density in controls and CKO. N=3, p = 0.187, unpaired student’s t-test. Data are presented as mean ± SD.

**Fig. EV5.**
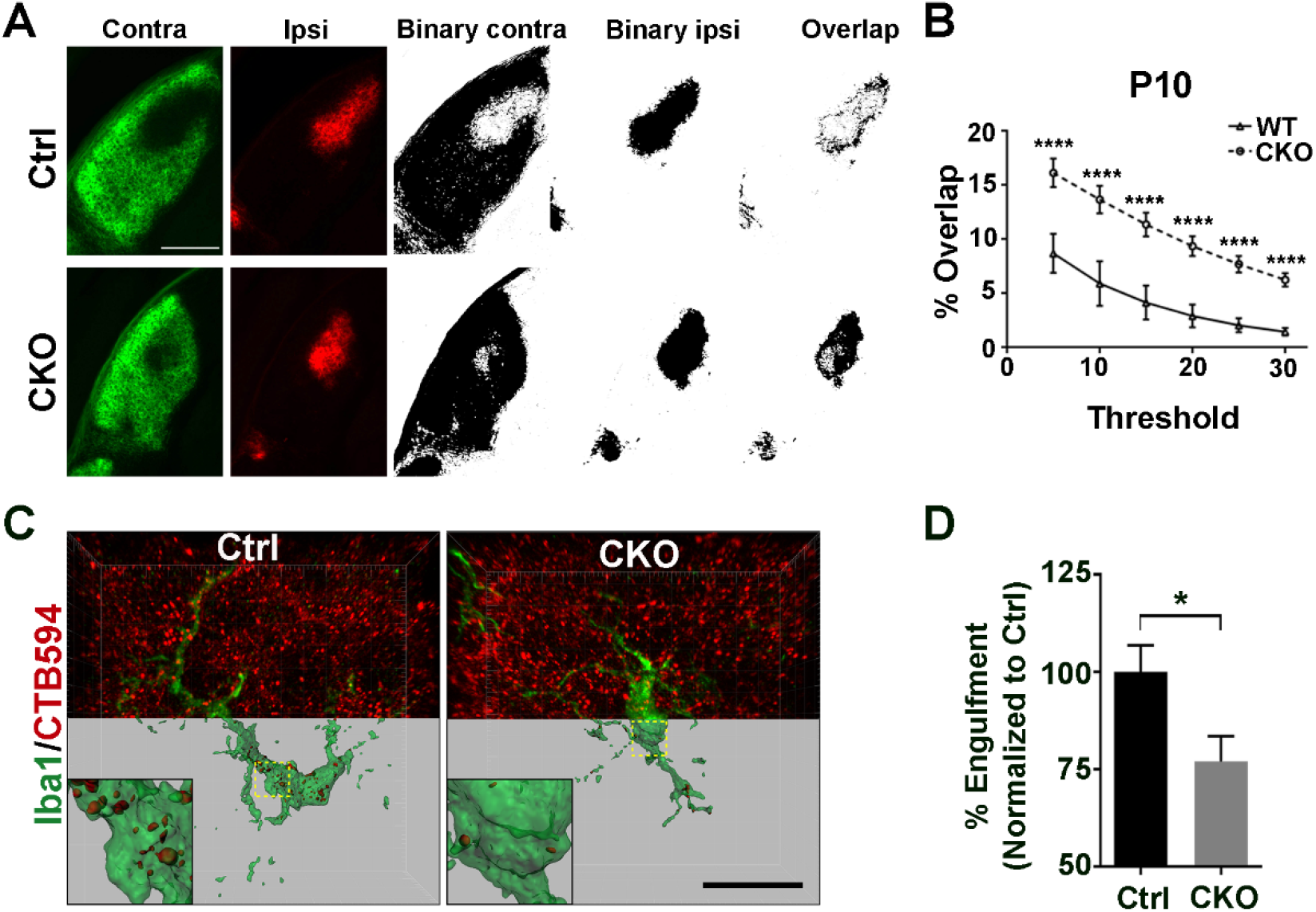
Deleting microglial *Gpr56* leads to impaired eye-specific segregation at P10, and reduced engulfment of RGC inputs by microglia. (A) The left two columns show contra- and ipsi-lateral RGC inputs labeled by CTB488 and CTB594, respectively. The middle two columns are binary images of contra- and ipsi-lateral LGN. The right images show an overlap between contralateral RGC inputs and ipsilateral inputs. (B) Quantification of the percentage of overlapped contra- and ipsi-lateral RGC inputs on multiple thresholds in CKO and controls at P10. N=4, ****p < 0.0001, two-way ANOVA with Bonferroni’s post hoc test. (C) Representative images and surface rendered microglia (green) from P5 dLGN of CKO or controls in which RGC inputs were labeled with CTB-594 (red). Scale bar, 20µm. (D) Quantification of the percentage of engulfed RGC inputs in controls and CKO microglia. More than 10 microglial cells are analyzed in each individual mouse brain. N = 4, p = 0.039, mean ± SEM.

## Appendix Figures

**Appendix Figure S1.**
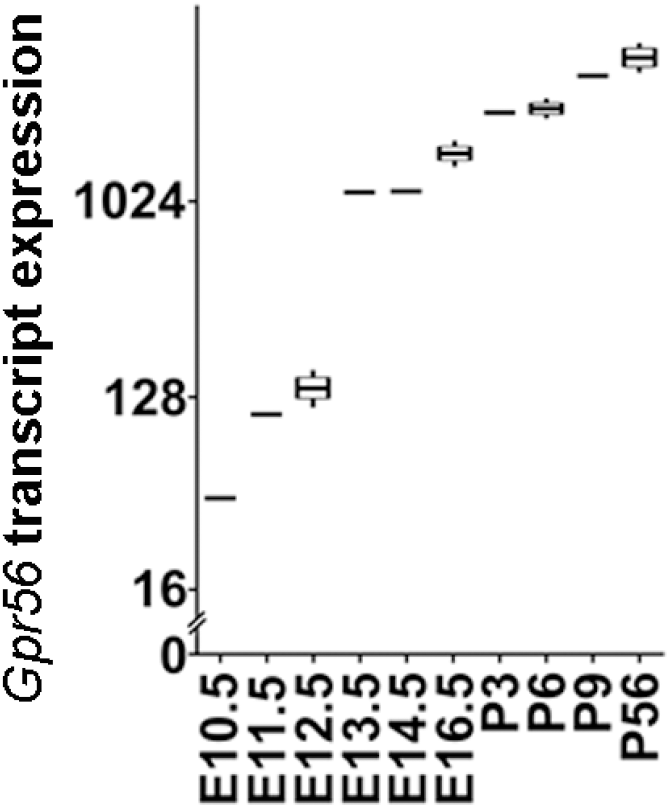
Microglial *Gpr56* transcript expression across different developmental stages. Data was obtained from the published RNAseq data (Matcovitch-Natan et al., 2016).

**Appendix Figure S2.**
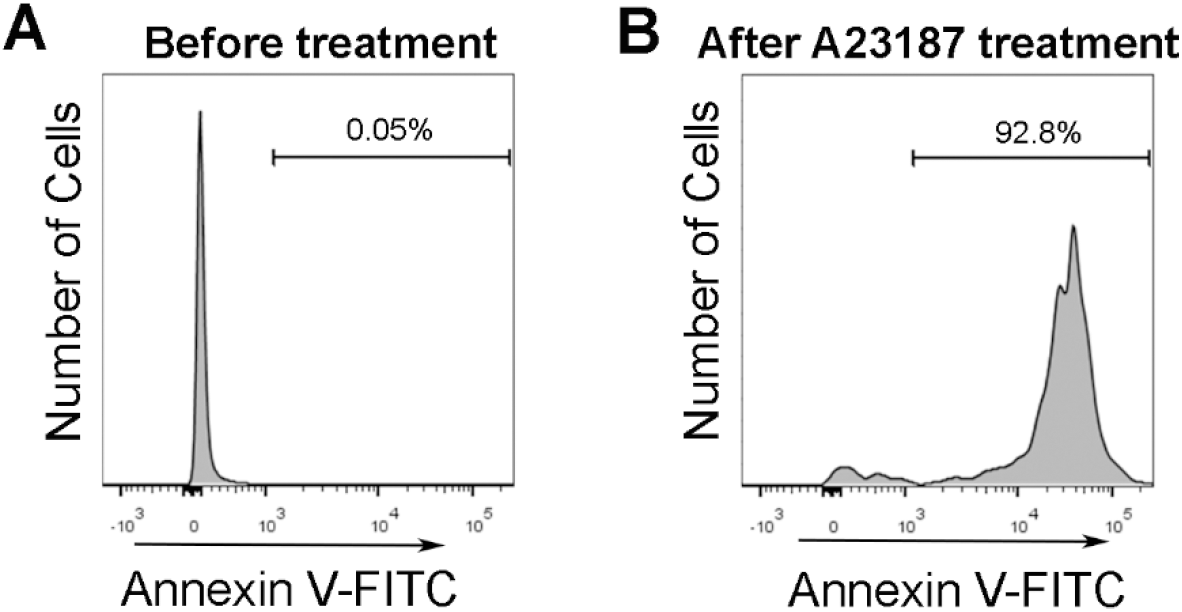
Exposure of PS on Ba/F3 cells after A23187 treatment. (A) Before treatment, Ba/F3 cells were barely detected by Annexin V, which binds to PS. (B) With treatment of calcium ionophore A23187, the majority of Ba/F3 became Annexin V positive, indicating Ba/F3 cells externalized PS to the outer leaflet of plasma membrane.

**Appendix Figure S3.**
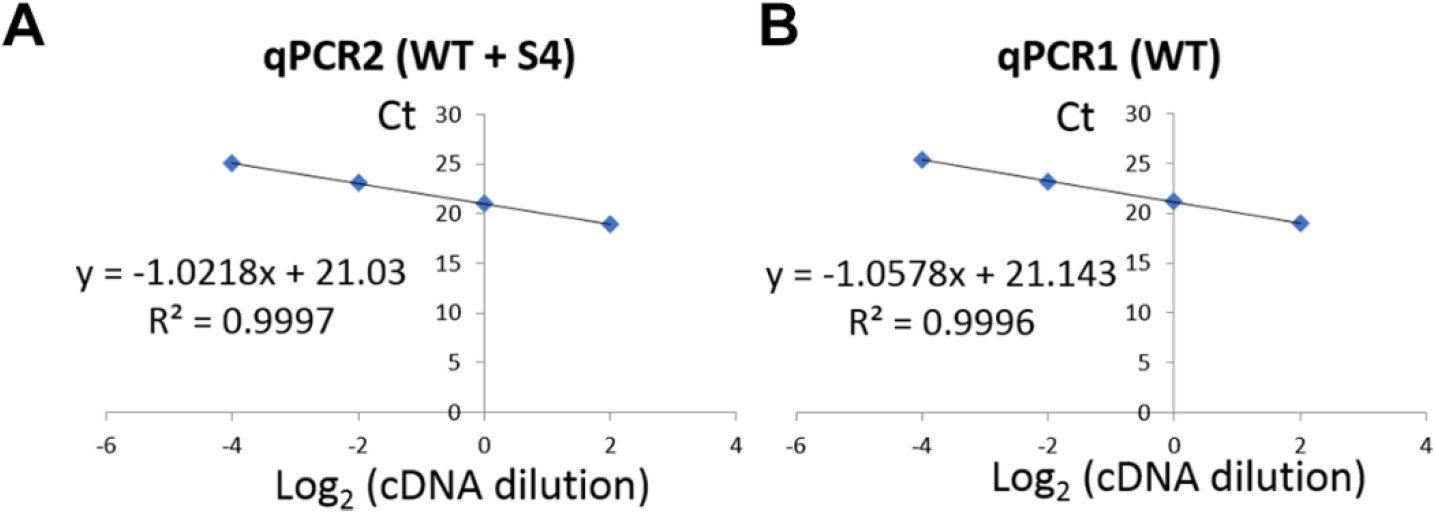
QPCR. (A) A standard curve for the qPCR2 reaction was performed using 1:2 serial dilution of microglial cDNA. (B) A standard curve for the qPCR1 reaction was performed.

**Appendix Figure S4.**
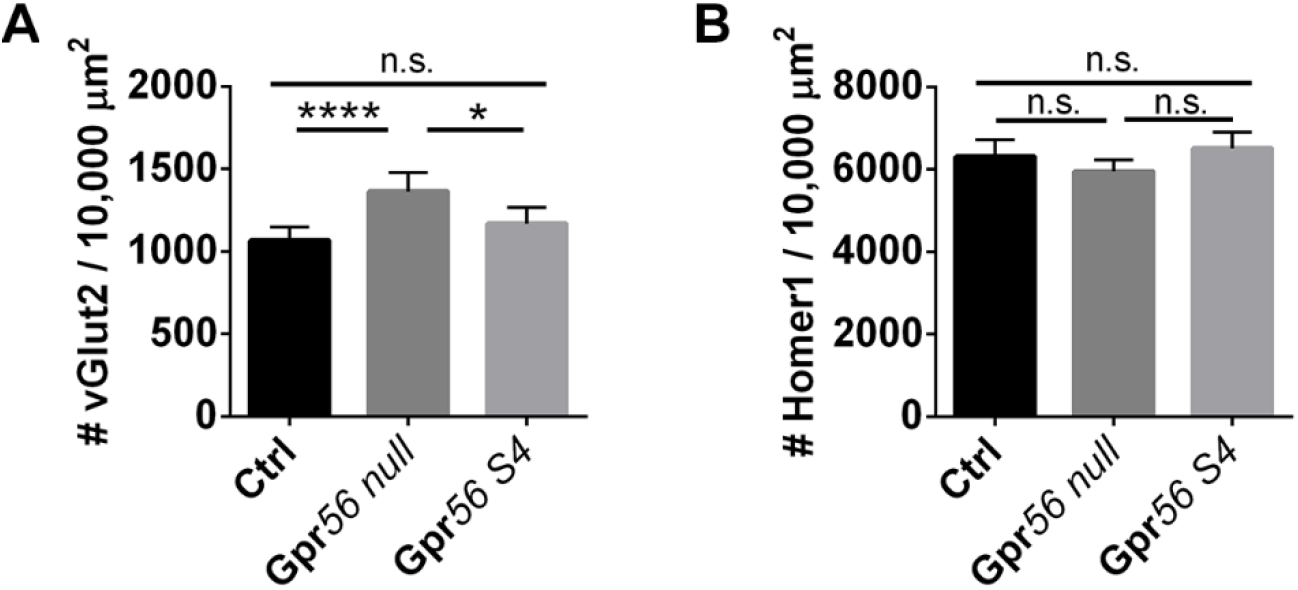
Quantification of vGlut2 presynapse and Homer1 postsynapse densities. (A) Increase vGlut2^+^ presynapse densities were found in *Gpr56 null* mice, not in *Gpr56 S4* mice. (B) No significant change of Homer1 postsynapse density was shown in *Gpr56 null* and *S4* mice. N=10 (Ctrl), N=6 (*Gpr56 null*), N=3 (*Gpr56 S4*). *p < 0.05, **** p < 0.0001, one-way ANOVA with Tukey’s post-hoc test.

**Appendix Figure S5.**
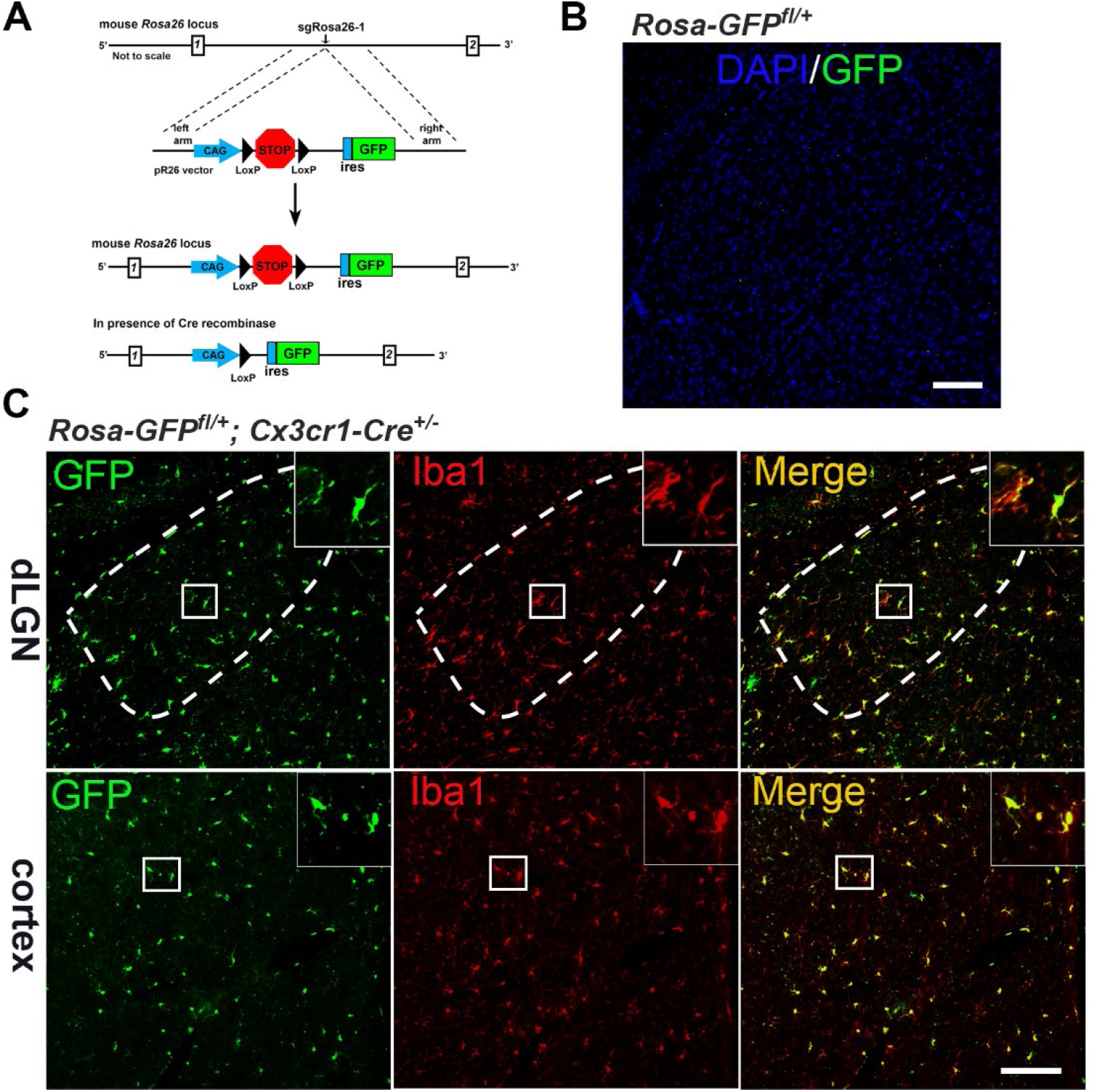
Restricted GFP expression in microglia driven by *Cx3cr1-Cre*. (A) A scheme showing the generation of *Rosa-GFP*^*fl*^ transgenic mice and *Rosa-GFP*^*fl/+*^; *Cx3cr1-Cre*^*+/-*^ mice. (B) GFP was barely observed in *Rosa-GFP*^*fl/+*^ mice. (C) GFP expression driven by *Cx3cr1-cre* was restricted in microglia in the dLGN and cortex. Scale bar, 100 µm.

**Appendix Figure S6.**
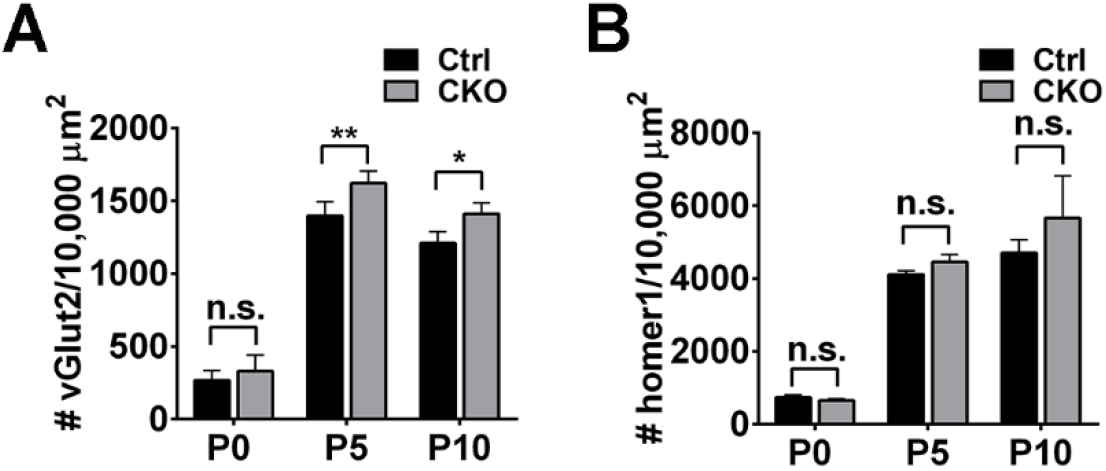
Quantification of vGlut2 presynapse and Homer1 postsynapse densities in early dLGN development. (A) Increase vGlut2^+^ presynapse densities were found in CKO mice at P5 and P10. (B) No significant change of Homer1 postsynapse density was seen in CKO mice. N = 3 (P0), N = 4 (P5), N=3 (P10). *p < 0.05, ** p < 0.01, two-way ANOVA followed by Bonferroni’s post hoc test.

**Appendix Figure S7.**
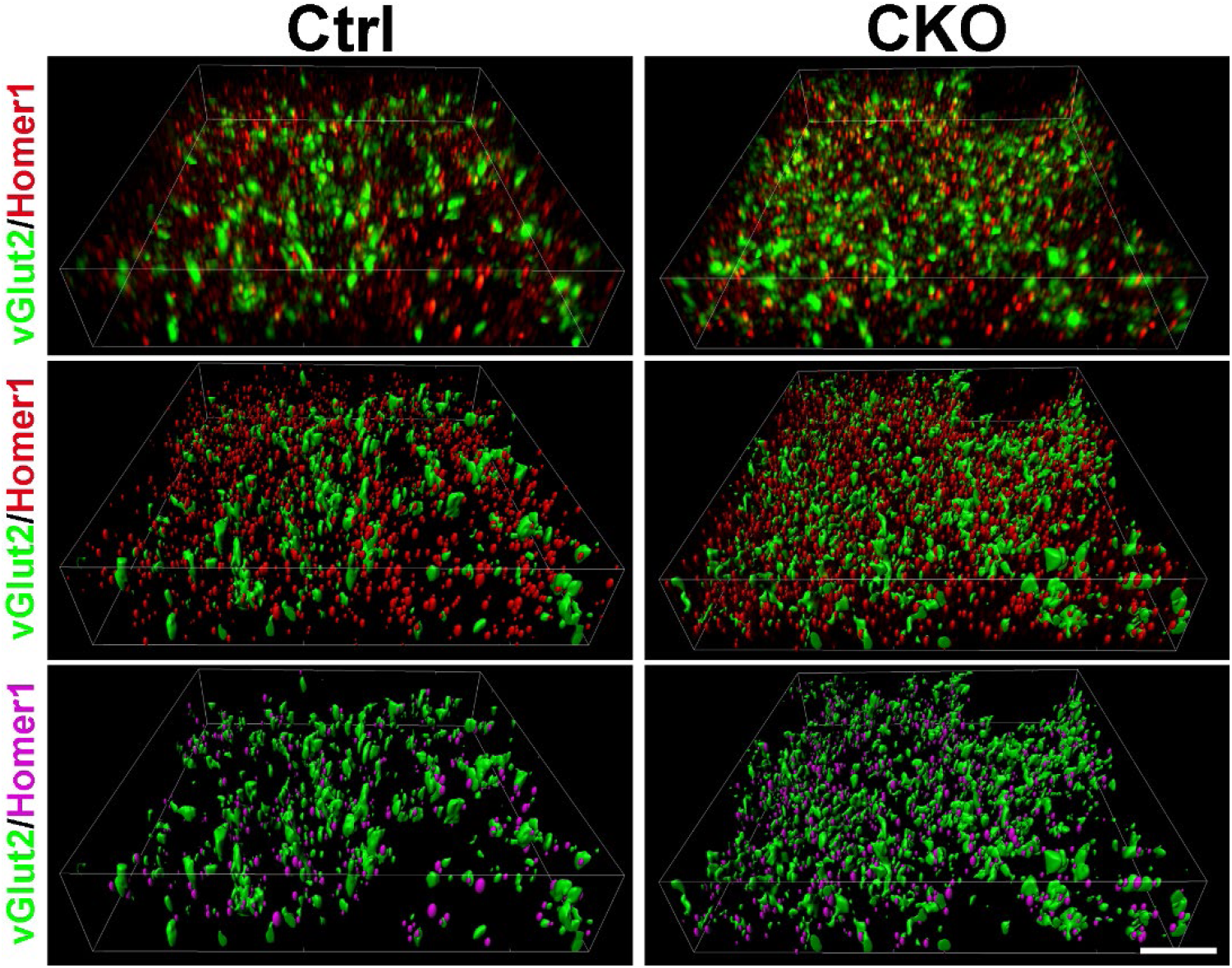
Retinogeniculate synapses in CKO and controls using SIM. Top panels show original images taken by SIM. Middle panels show 3D rendered images after processing in Imaris. Bottom panels show Homer1 spots (magenta) that are within the distance of 300nm from vGlut2 surface (green). Scale bar, 5 µm.

**Appendix Figure S8.**
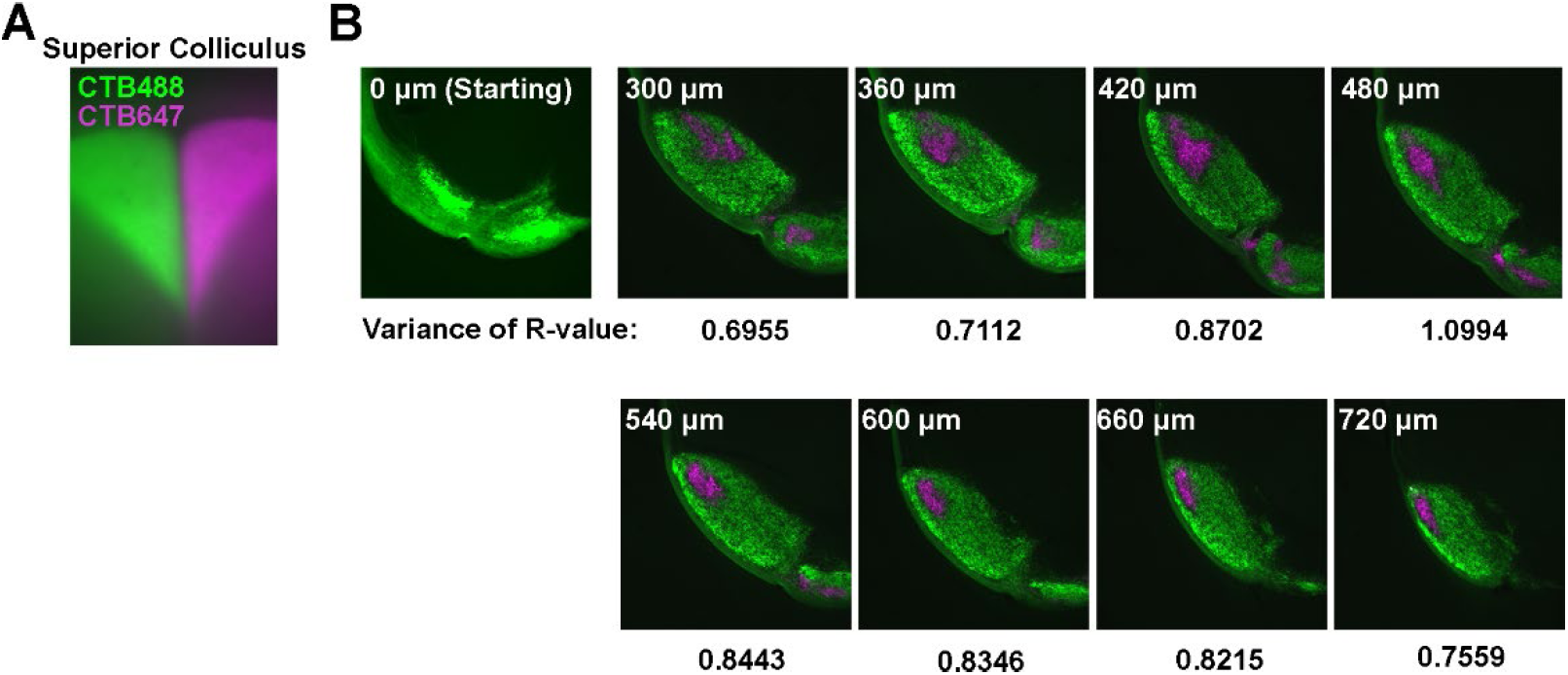
The variance of R-value varies from cranial to caudal LGN. (A) Images of superior colliculus after 24 hours anterograde labeling of RGC by CTB488 and CTB647 at P30, indicating efficient RGC labeling by CTB. (B) Variance of R-values varies from caudal LGN to cranial LGN.

**Appendix Figure S9.**
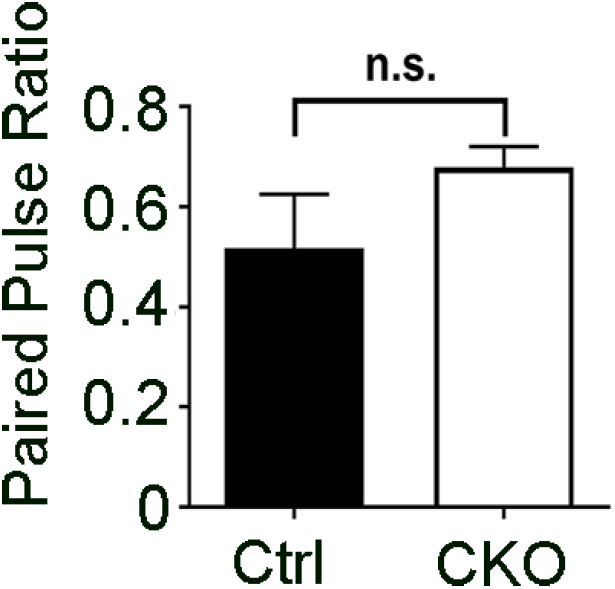
*Gpr56* CKO mice show no significant difference in paired pulse depression. Paired pulse depression was recorded on dLGN slice from P28-P34 mice. Given that optic inputs usually demonstrate paired pulse depression, and cortical inputs show paired pulse facilitation, this data indicates that the optic tracts and not cortical inputs were stimulated. N = 14 (Ctrl), 23 (CKO) cells from 5, 7 mice. P = 0.694 by Student’s t-test.

**Movie EV1. 3D reconstruction of a SIM image of synapses.**

(A) A representative image of vGlut2^+^ pre-synapses and Homer1^+^ post-synapses taken by SIM. (B) 3D reconstruction of the image in (A). (C) Purple spots represent Homer1^+^ post-synapses within the distance of 300nm from vGlut2^+^ (green) surface.

## References

Ackerman, S.D., Garcia, C., Piao, X., Gutmann, D.H., and Monk, K.R. (2015). The adhesion GPCR Gpr56 regulates oligodendrocyte development via interactions with Gα 12/13 and RhoA. Nat Commun 6, 6122.

Araç, D., Boucard, A.A., Bolliger, M.F., Nguyen, J., Soltis, S.M., Sudhof, T.C., and Brunger, A.T. (2012). A novel evolutionarily conserved domain of cell-adhesion GPCRs mediates autoproteolysis. EMBO J 31, 1364–1378.

Bennett, F.C., Bennett, M.L., Yaqoob, F., Mulinyawe, S.B., Grant, G.A., Hayden Gephart, M., Plowey, E.D., and Barres, B.A. (2018). A Combination of Ontogeny and CNS Environment Establishes Microglial Identity. Neuron 98, 1170–1183 e1178.

Bennett, M.L., Bennett, F.C., Liddelow, S.A., Ajami, B., Zamanian, J.L., Fernhoff, N.B., Mulinyawe, S.B., Bohlen, C.J., Adil, A., Tucker, A., et al. (2016). New tools for studying microglia in the mouse and human CNS. Proc Natl Acad Sci USA 113, E1738–1746.

Bevers, E.M., and Williamson, P.L. (2016). Getting to the Outer Leaflet: Physiology of Phosphatidylserine Exposure at the Plasma Membrane. Physiol Rev 96, 605–645.

Bialas, A.R., Presumey, J., Das, A., van der Poel, C.E., Lapchak, P.H., Mesin, L., Victora, G., Tsokos, G.C., Mawrin, C., Herbst, R., and Carroll, M.C. (2017). Microglia-dependent synapse loss in type I interferon-mediated lupus. Nature 546, 539–543.

Bohlen, C.J., Bennett, F.C., Tucker, A.F., Collins, H.Y., Mulinyawe, S.B., and Barres, B.A. (2017). Diverse Requirements for Microglial Survival, Specification, and Function Revealed by Defined-Medium Cultures. Neuron 94, 759–773 e758.

Braunschweig, U., Gueroussov, S., Plocik, A.M., Graveley, B.R., and Blencowe, B.J. (2013). Dynamic integration of splicing within gene regulatory pathways. Cell 152, 1252–1269.

Cermak, N., Olcum, S., Delgado, F.F., Wasserman, S.C., Payer, K.R., M, A.M., Knudsen, S.M., Kimmerling, R.J., Stevens, M.M., Kikuchi, Y., et al. (2016). High-throughput measurement of single-cell growth rates using serial microfluidic mass sensor arrays. Nat Biotechnol 34, 1052–1059.

Chen, C., and Regehr, W.G. (2000). Developmental remodeling of the retinogeniculate synapse. Neuron 28, 955–966.

Chu, V.T., Weber, T., Graf, R., Sommermann, T., Petsch, K., Sack, U., Volchkov, P., Rajewsky, K., and Kuhn, R. (2016). Efficient generation of Rosa26 knock-in mice using CRISPR/Cas9 in C57BL/6 zygotes. BMC Biotechnol 16, 4.

Chung, W.S., Clarke, L.E., Wang, G.X., Stafford, B.K., Sher, A., Chakraborty, C., Joung, J., Foo, L.C., Thompson, A., Chen, C., et al. (2013). Astrocytes mediate synapse elimination through MEGF10 and MERTK pathways. Nature 504, 394–400.

Cronk, J.C., Filiano, A.J., Louveau, A., Marin, I., Marsh, R., Ji, E., Goldman, D.H., Smirnov, I., Geraci, N., Acton, S., et al. (2018). Peripherally derived macrophages can engraft the brain independent of irradiation and maintain an identity distinct from microglia. J Exp Med 215, 1627–1647.

Dani, A., Huang, B., Bergan, J., Dulac, C., and Zhuang, X. (2010). Superresolution imaging of chemical synapses in the brain. Neuron 68, 843–856.

Djurisic, M., Brott, B.K., Saw, N.L., Shamloo, M., and Shatz, C.J. (2019). Activity-dependent modulation of hippocampal synaptic plasticity via PirB and endocannabinoids. Mol Psychiatry 24, 1206–1219.

Erturk, A., Wang, Y., and Sheng, M. (2014). Local pruning of dendrites and spines by caspase-3-dependent and proteasome-limited mechanisms. J Neurosci 34, 1672–1688.

Etxeberria, A., Hokanson, K.C., Dao, D.Q., Mayoral, S.R., Mei, F., Redmond, S.A., Ullian, E.M., and Chan, J.R. (2016). Dynamic Modulation of Myelination in Response to Visual Stimuli Alters Optic Nerve Conduction Velocity. J Neurosci 36, 6937–6948.

Feng, W., Yasumura, D., Matthes, M.T., LaVail, M.M., and Vollrath, D. (2002). Mertk triggers uptake of photoreceptor outer segments during phagocytosis by cultured retinal pigment epithelial cells. J Biol Chem 277, 17016–17022.

Fitzpatrick, D., Usrey, W.M., Schofield, B.R., and Einstein, G. (1994). The sublaminar organization of corticogeniculate neurons in layer 6 of macaque striate cortex. Vis Neurosci 11, 307–315.

Folts, C.J., Giera, S., Li, T., and Piao, X. (2019). Adhesion G Protein-Coupled Receptors as Drug Targets for Neurological Diseases. Trends Pharmacol Sci 40, 278–293.

Frank, M.G., Wieseler-Frank, J.L., Watkins, L.R., and Maier, S.F. (2006). Rapid isolation of highly enriched and quiescent microglia from adult rat hippocampus: immunophenotypic and functional characteristics. J Neurosci Methods 151, 121–130.

Fujiyama, F., Hioki, H., Tomioka, R., Taki, K., Tamamaki, N., Nomura, S., Okamoto, K., and Kaneko, T. (2003). Changes of immunocytochemical localization of vesicular glutamate transporters in the rat visual system after the retinofugal denervation. J Comp Neurol 465, 234–249.

Fujiyama, F., Kuramoto, E., Okamoto, K., Hioki, H., Furuta, T., Zhou, L., Nomura, S., and Kaneko, T. (2004). Presynaptic localization of an AMPA-type glutamate receptor in corticostriatal and thalamostriatal axon terminals. Eur J Neurosci 20, 3322–3330.

Giera, S., Deng, Y., Luo, R., Ackerman, S.D., Mogha, A., Monk, K.R., Ying, Y., Jeong, S.J., Makinodan, M., Bialas, A.R., et al. (2015). The adhesion G protein-coupled receptor GPR56 is a cell-autonomous regulator of oligodendrocyte development. Nat Commun 6, 6121.

Giera, S., Luo, R., Ying, Y., Ackerman, S.D., Jeong, S.J., Stoveken, H.M., Folts, C.J., Welsh, C.A., Tall, G.G., Stevens, B., et al. (2018). Microglial transglutaminase-2 drives myelination and myelin repair via GPR56/ADGRG1 in oligodendrocyte precursor cells. Elife 7, e33385.

Gosselin, D., Link, V.M., Romanoski, C.E., Fonseca, G.J., Eichenfield, D.Z., Spann, N.J., Stender, J.D., Chun, H.B., Garner, H., Geissmann, F., and Glass, C.K. (2014). Environment drives selection and function of enhancers controlling tissue-specific macrophage identities. Cell 159, 1327–1340.

Gosselin, D., Skola, D., Coufal, N.G., Holtman, I.R., Schlachetzki, J.C.M., Sajti, E., Jaeger, B.N., O’Connor, C., Fitzpatrick, C., Pasillas, M.P., et al. (2017). An environment-dependent transcriptional network specifies human microglia identity. Science 356.

Gyorffy, B.A., Kun, J., Torok, G., Bulyaki, E., Borhegyi, Z., Gulyassy, P., Kis, V., Szocsics, P., Micsonai, A., Matko, J., et al. (2018). Local apoptotic-like mechanisms underlie complement-mediated synaptic pruning. Proc Natl Acad Sci USA 115, 6303–6308.

Hagemeyer, N., Hanft, K.M., Akriditou, M.A., Unger, N., Park, E.S., Stanley, E.R., Staszewski, O., Dimou, L., and Prinz, M. (2017). Microglia contribute to normal myelinogenesis and to oligodendrocyte progenitor maintenance during adulthood. Acta Neuropathol 134, 441–458.

Hamann, J., Aust, G., Arac, D., Engel, F.B., Formstone, C., Fredriksson, R., Hall, R.A., Harty, B.L., Kirchhoff, C., Knapp, B., et al. (2015). International Union of Basic and Clinical Pharmacology. XCIV. Adhesion G protein-coupled receptors. Pharmacol Rev 67, 338–367.

Hanshaw, R.G., and Smith, B.D. (2005). New reagents for phosphatidylserine recognition and detection of apoptosis. Bioorg Med Chem 13, 5035–5042.

Hong, S., Beja-Glasser, V.F., Nfonoyim, B.M., Frouin, A., Li, S., Ramakrishnan, S., Merry, K.M., Shi, Q., Rosenthal, A., Barres, B.A., et al. (2016). Complement and microglia mediate early synapse loss in Alzheimer mouse models. Science 352, 712–716.

Hong, S., Wilton, D.K., Stevens, B., and Richardson, D.S. (2017). Structured Illumination Microscopy for the Investigation of Synaptic Structure and Function. Methods Mol Biol 1538, 155–167.

Ingolfsson, H.I., Melo, M.N., van Eerden, F.J., Arnarez, C., Lopez, C.A., Wassenaar, T.A., Periole, X., de Vries, A.H., Tieleman, D.P., and Marrink, S.J. (2014). Lipid organization of the plasma membrane. J Am Chem Soc 136, 14554–14559.

Jeong, S.J., Li, S., Luo, R., Strokes, N., and Piao, X. (2012a). Loss of Col3a1, the gene for Ehlers-Danlos syndrome type IV, results in neocortical dyslamination. PLoS One 7, e29767.

Jeong, S.J., Luo, R., Li, S., Strokes, N., and Piao, X. (2012b). Characterization of G protein-coupled receptor 56 protein expression in the mouse developing neocortex. J Comp Neurol 520, 2930–2940.

Kim, J.E., Han, J.M., Park, C.R., Shin, K.J., Ahn, C., Seong, J.Y., and Hwang, J.I. (2010). Splicing variants of the orphan G-protein-coupled receptor GPR56 regulate the activity of transcription factors associated with tumorigenesis. J Cancer Res Clin Oncol 136, 47–53.

Koopman, G., Reutelingsperger, C.P., Kuijten, G.A., Keehnen, R.M., Pals, S.T., and van Oers, M.H. (1994). Annexin V for flow cytometric detection of phosphatidylserine expression on B cells undergoing apoptosis. Blood 84, 1415–1420.

Koulov, A.V., Stucker, K.A., Lakshmi, C., Robinson, J.P., and Smith, B.D. (2003). Detection of apoptotic cells using a synthetic fluorescent sensor for membrane surfaces that contain phosphatidylserine. Cell Death Differ 10, 1357–1359.

Land, P.W., Kyonka, E., and Shamalla-Hannah, L. (2004). Vesicular glutamate transporters in the lateral geniculate nucleus: expression of VGLUT2 by retinal terminals. Brain Res 996, 251–254.

Langenhan, T., Piao, X., and Monk, K.R. (2016). Adhesion G protein-coupled receptors in nervous system development and disease. Nat Rev Neurosci 17, 550–561.

Lee, H., Brott, B.K., Kirkby, L.A., Adelson, J.D., Cheng, S., Feller, M.B., Datwani, A., and Shatz, C.J. (2014). Synapse elimination and learning rules co-regulated by MHC class I H2-Db. Nature 509, 195–200.

Lehrman, E.K., Wilton, D.K., Litvina, E.Y., Welsh, C.A., Chang, S.T., Frouin, A., Walker, A.J., Heller, M.D., Umemori, H., Chen, C., and Stevens, B. (2018). CD47 Protects Synapses from Excess Microglia-Mediated Pruning during Development. Neuron 100, 120-134.e126.

Lemke, G. (2013). Biology of the TAM receptors. Cold Spring Harb Perspect Biol 5, a009076.

Li, S., Jin, Z., Koirala, S., Bu, L., Xu, L., Hynes, R.O., Walsh, C.A., Corfas, G., and Piao, X. (2008). GPR56 regulates pial basement membrane integrity and cortical lamination. J Neurosci 28, 5817–5826.

Luo, R., Jeong, S.J., Jin, Z., Strokes, N., Li, S., and Piao, X. (2011). G protein-coupled receptor 56 and collagen III, a receptor-ligand pair, regulates cortical development and lamination. Proc Natl Acad Sci USA 108, 12925–12930.

Luo, R., Jin, Z., Deng, Y., Strokes, N., and Piao, X. (2012). Disease-associated mutations prevent GPR56-collagen III interaction. PLoS One 7, e29818.

Mark, M.R., Chen, J., Hammonds, R.G., Sadick, M., and Godowsk, P.J. (1996). Characterization of Gas6, a member of the superfamily of G domain-containing proteins, as a ligand for Rse and Axl. J Biol Chem 271, 9785–9789.

Matcovitch-Natan, O., Winter, D.R., Giladi, A., Vargas Aguilar, S., Spinrad, A., Sarrazin, S., Ben-Yehuda, H., David, E., Zelada Gonzalez, F., Perrin, P., et al. (2016). Microglia development follows a stepwise program to regulate brain homeostasis. Science 353, aad8670.

Mattson, M.P., Keller, J.N., and Begley, J.G. (1998). Evidence for synaptic apoptosis. Exp Neurol 153, 35–48.

Mayoral, S.R., Etxeberria, A., Shen, Y.A., and Chan, J.R. (2018). Initiation of CNS Myelination in the Optic Nerve Is Dependent on Axon Caliber. Cell Rep 25, 544-550.e543.

Mazaheri, F., Breus, O., Durdu, S., Haas, P., Wittbrodt, J., Gilmour, D., and Peri, F. (2014). Distinct roles for BAI1 and TIM-4 in the engulfment of dying neurons by microglia. Nat Commun 5, 4046.

McLaughlin, S., and Murray, D. (2005). Plasma membrane phosphoinositide organization by protein electrostatics. Nature 438, 605–611.

Miyamoto, A., Wake, H., Ishikawa, A.W., Eto, K., Shibata, K., Murakoshi, H., Koizumi, S., Moorhouse, A.J., Yoshimura, Y., and Nabekura, J. (2016). Microglia contact induces synapse formation in developing somatosensory cortex. Nat Commun 7, 12540.

Miyanishi, M., Tada, K., Koike, M., Uchiyama, Y., Kitamura, T., and Nagata, S. (2007). Identification of Tim4 as a phosphatidylserine receptor. Nature 450, 435–439.

Nadal-Nicolas, F.M., Jimenez-Lopez, M., Sobrado-Calvo, P., Nieto-Lopez, L., Canovas-Martinez, I., Salinas-Navarro, M., Vidal-Sanz, M., and Agudo, M. (2009). Brn3a as a marker of retinal ganglion cells: qualitative and quantitative time course studies in naive and optic nerve-injured retinas. Invest Ophthalmol Vis Sci 50, 3860–3868.

Nagata, K., Ohashi, K., Nakano, T., Arita, H., Zong, C., Hanafusa, H., and Mizuno, K. (1996). Identification of the product of growth arrest-specific gene 6 as a common ligand for Axl, Sky, and Mer receptor tyrosine kinases. J Biol Chem 271, 30022–30027.

Nagata, S., Suzuki, J., Segawa, K., and Fujii, T. (2016). Exposure of phosphatidylserine on the cell surface. Cell Death Differ 23, 952–961.

Neher, J.J., Emmrich, J.V., Fricker, M., Mander, P.K., Thery, C., and Brown, G.C. (2013). Phagocytosis executes delayed neuronal death after focal brain ischemia. Proc Natl Acad Sci USA 110, E4098–4107.

Ohashi, K., Nagata, K., Toshima, J., Nakano, T., Arita, H., Tsuda, H., Suzuki, K., and Mizuno, K. (1995). Stimulation of sky receptor tyrosine kinase by the product of growth arrest-specific gene 6. J Biol Chem 270, 22681–22684.

Paolicelli, R.C., Bolasco, G., Pagani, F., Maggi, L., Scianni, M., Panzanelli, P., Giustetto, M., Ferreira, T.A., Guiducci, E., Dumas, L., et al. (2011). Synaptic pruning by microglia is necessary for normal brain development. Science 333, 1456–1458.

Paradies, G., Paradies, V., De Benedictis, V., Ruggiero, F.M., and Petrosillo, G. (2014). Functional role of cardiolipin in mitochondrial bioenergetics. Biochim Biophys Acta 1837, 408–417.

Park, D., Tosello-Trampont, A.C., Elliott, M.R., Lu, M., Haney, L.B., Ma, Z., Klibanov, A.L., Mandell, J.W., and Ravichandran, K.S. (2007). BAI1 is an engulfment receptor for apoptotic cells upstream of the ELMO/Dock180/Rac module. Nature 450, 430–434.

Park, S.Y., Jung, M.Y., Kim, H.J., Lee, S.J., Kim, S.Y., Lee, B.H., Kwon, T.H., Park, R.W., and Kim, I.S. (2008). Rapid cell corpse clearance by stabilin-2, a membrane phosphatidylserine receptor. Cell Death Differ 15, 192–201.

Parkhurst, C.N., Yang, G., Ninan, I., Savas, J.N., Yates III, J.R., Lafaille, J.J., Hempstead, B.L., Littman, D.R., and Gan, W.B. (2013). Microglia promote learning-dependent synapse formation through brain-derived neurotrophic factor. Cell 155, 1596–1609.

Piao, X., Chang, B.S., Bodell, A., Woods, K., Benzeev, B., Topcu, M., Guerrini, R., Goldberg-Stern, H., Sztriha, L., Dobyns, W.B., et al. (2005). Genotype-phenotype analysis of human frontoparietal polymicrogyria syndromes. Ann Neurol 58, 680–687.

Piao, X., Hill, R.S., Bodell, A., Chang, B.S., Basel-Vanagaite, L., Straussberg, R., Dobyns, W.B., Qasrawi, B., Winter, R.M., Innes, A.M., et al. (2004). G protein-coupled receptor-dependent development of human frontal cortex. Science 303, 2033–2036.

Reemst, K., Noctor, S.C., Lucassen, P.J., and Hol, E.M. (2016). The Indispensable Roles of Microglia and Astrocytes during Brain Development. Front Hum Neurosci 10, 566.

Salzman, G.S., Ackerman, S.D., Ding, C., Koide, A., Leon, K., Luo, R., Stoveken, H.M., Fernandez, C.G., Tall, G.G., Piao, X., et al. (2016). Structural Basis for Regulation of GPR56/ADGRG1 by Its Alternatively Spliced Extracellular Domains. Neuron 91, 1292–1304.

Sato, K. (2015). Effects of Microglia on Neurogenesis. Glia 63, 1394–1405.

Schafer, D.P., Lehrman, E.K., Kautzman, A.G., Koyama, R., Mardinly, A.R., Yamasaki, R., Ransohoff, R.M., Greenberg, M.E., Barres, B.A., and Stevens, B. (2012). Microglia sculpt postnatal neural circuits in an activity and complement-dependent manner. Neuron 74, 691–705.

Schwenk, F., Baron, U., and Rajewsky, K. (1995). A cre-transgenic mouse strain for the ubiquitous deletion of loxP-flanked gene segments including deletion in germ cells. Nucleic Acids Res 23, 5080–5081.

Segawa, K., and Nagata, S. (2015). An Apoptotic ‘Eat Me’ Signal: Phosphatidylserine Exposure. Trends Cell Biol 25, 639–650.

Segawa, K., Suzuki, J., and Nagata, S. (2011). Constitutive exposure of phosphatidylserine on viable cells. Proc Natl Acad Sci USA 108, 19246–19251.

Sekar, A., Bialas, A.R., de Rivera, H., Davis, A., Hammond, T.R., Kamitaki, N., Tooley, K., Presumey, J., Baum, M., Van Doren, V., et al. (2016). Schizophrenia risk from complex variation of complement component 4. Nature 530, 177–183.

Sellgren, C.M., Gracias, J., Watmuff, B., Biag, J.D., Thanos, J.M., Whittredge, P.B., Fu, T., Worringer, K., Brown, H.E., Wang, J., et al. (2019). Increased synapse elimination by microglia in schizophrenia patient-derived models of synaptic pruning. Nat Neurosci 22, 374–385.

Shatz, C.J., and Kirkwood, P.A. (1984). Prenatal development of functional connections in the cat’s retinogeniculate pathway. J Neurosci 4, 1378–1397.

Sherman, S.M., and Guillery, R.W. (1996). Functional organization of thalamocortical relays. J Neurophysiol 76, 1367–1395.

Sherman, S.M., and Guillery, R.W. (1998). On the actions that one nerve cell can have on another: distinguishing “drivers” from “modulators”. Proc Natl Acad Sci USA 95, 7121–7126.

Singer, K., Luo, R., Jeong, S.J., and Piao, X. (2013). GPR56 and the developing cerebral cortex: cells, matrix, and neuronal migration. Mol Neurobiol 47, 186–196.

Smith, B.A., Gammon, S.T., Xiao, S., Wang, W., Chapman, S., McDermott, R., Suckow, M.A., Johnson, J.R., Piwnica-Worms, D., Gokel, G.W., et al. (2011). In vivo optical imaging of acute cell death using a near-infrared fluorescent zinc-dipicolylamine probe. Mol Pharm 8, 583–590.

Sretavan, D., and Shatz, C.J. (1984). Prenatal development of individual retinogeniculate axons during the period of segregation. Nature 308, 845–848.

Stevens, B., Allen, N.J., Vazquez, L.E., Howell, G.R., Christopherson, K.S., Nouri, N., Micheva, K.D., Mehalow, A.K., Huberman, A.D., Stafford, B., et al. (2007). The classical complement cascade mediates CNS synapse elimination. Cell 131, 1164–1178.

Stitt, T.N., Conn, G., Gore, M., Lai, C., Bruno, J., Radziejewski, C., Mattsson, K., Fisher, J., Gies, D.R., Jones, P.F., and et al. (1995). The anticoagulation factor protein S and its relative, Gas6, are ligands for the Tyro 3/Axl family of receptor tyrosine kinases. Cell 80, 661–670.

Suh, J., Rivest, A.J., Nakashiba, T., Tominaga, T., and Tonegawa, S. (2011). Entorhinal cortex layer III input to the hippocampus is crucial for temporal association memory. Science 334, 1415–1420.

Suzuki, J., Umeda, M., Sims, P.J., and Nagata, S. (2010). Calcium-dependent phospholipid scrambling by TMEM16F. Nature 468, 834–838.

Swanson, L.W., Wyss, J.M., and Cowan, W.M. (1978). An autoradiographic study of the organization of intrahippocampal association pathways in the rat. J Comp Neurol 181, 681–715.

Tang, G., Gudsnuk, K., Kuo, S.H., Cotrina, M.L., Rosoklija, G., Sosunov, A., Sonders, M.S., Kanter, E., Castagna, C., Yamamoto, A., et al. (2014). Loss of mTOR-dependent macroautophagy causes autistic-like synaptic pruning deficits. Neuron 83, 1131–1143.

Thomson, A.M. (2010). Neocortical layer 6, a review. Front Neuroanat 4, 13.

Tiriac, A., Smith, B.E., and Feller, M.B. (2018). Light Prior to Eye Opening Promotes Retinal Waves and Eye-Specific Segregation. Neuron 100, 1059-1065.e1054.

Torborg, C.L., and Feller, M.B. (2004). Unbiased analysis of bulk axonal segregation patterns. J Neurosci Methods 135, 17–26.

Tufail, Y., Cook, D., Fourgeaud, L., Powers, C.J., Merten, K., Clark, C.L., Hoffman, E., Ngo, A., Sekiguchi, K.J., O’Shea, C.C., et al. (2017). Phosphatidylserine Exposure Controls Viral Innate Immune Responses by Microglia. Neuron 93, 574–586 e578.

Turner, J.P., and Salt, T.E. (1998). Characterization of sensory and corticothalamic excitatory inputs to rat thalamocortical neurones *in vitro*. J Physiol 510 (Pt 3), 829-843.

Vainchtein, I.D., Chin, G., Cho, F.S., Kelley, K.W., Miller, J.G., Chien, E.C., Liddelow, S.A., Nguyen, P.T., Nakao-Inoue, H., Dorman, L.C., et al. (2018). Astrocyte-derived interleukin-33 promotes microglial synapse engulfment and neural circuit development. Science 359, 1269–1273.

Vasek, M.J., Garber, C., Dorsey, D., Durrant, D.M., Bollman, B., Soung, A., Yu, J., Perez-Torres, C., Frouin, A., Wilton, D.K., et al. (2016). A complement-microglial axis drives synapse loss during virus-induced memory impairment. Nature 534, 538–543.

Yang, L., Chen, G., Mohanty, S., Scott, G., Fazal, F., Rahman, A., Begum, S., Hynes, R.O., and Xu, L. (2011). GPR56 Regulates VEGF production and angiogenesis during melanoma progression. Cancer Res 71, 5558–5568.

Yona, S., Kim, K.W., Wolf, Y., Mildner, A., Varol, D., Breker, M., Strauss-Ayali, D., Viukov, S., Guilliams, M., Misharin, A., et al. (2013). Fate mapping reveals origins and dynamics of monocytes and tissue macrophages under homeostasis. Immunity 38, 79–91.

Zachowski, A., Henry, J.P., and Devaux, P.F. (1989). Control of transmembrane lipid asymmetry in chromaffin granules by an ATP-dependent protein. Nature 340, 75–76.

Zhang, Y., Chen, K., Sloan, S.A., Bennett, M.L., Scholze, A.R., O’Keeffe, S., Phatnani, H.P., Guarnieri, P., Caneda, C., Ruderisch, N., et al. (2014). An RNA-sequencing transcriptome and splicing database of glia, neurons, and vascular cells of the cerebral cortex. J Neurosci 34, 11929–11947.

